# Application of a bioinformatic pipeline to RNA-seq data identifies novel virus-like sequence in human blood

**DOI:** 10.1101/2021.01.27.428546

**Authors:** Marko Melnick, Patrick Gonzales, Thomas J. LaRocca, Robin D. Dowell, Yuping Song, Joanne Wuu, Michael Benatar, Björn Oskarsson, Leonard Petrucelli, Christopher D. Link, Mercedes Prudencio

## Abstract

Numerous reports have suggested that infectious agents could play a role in neurodegenerative diseases, but specific etiological agents have not been convincingly demonstrated. To search for candidate agents in an unbiased fashion, we have developed a bioinformatic pipeline that identifies microbial sequences in mammalian RNA-seq data, including sequences with no significant nucleotide similarity hits in GenBank. Effectiveness of the pipeline was tested using publicly available RNA-seq data. We then applied this pipeline to a novel RNA-seq dataset generated from a cohort of 120 samples from amyotrophic lateral sclerosis (ALS) patients and controls, and identified sequences corresponding to known bacteria and viruses, as well as novel virus-like sequences. The presence of these novel virus-like sequences, which were identified in subsets of both patients and controls, were confirmed by quantitative RT-PCR. We believe this pipeline will be a useful tool for the identification of potential etiological agents in the many RNA-seq data sets currently being generated.

## INTRODUCTION

### Background of organisms in neurodegeneration

Infection has been proposed to play a role in multiple neurodegenerative diseases^1^, including amyotrophic lateral sclerosis (ALS)^2^. ALS is the most common motor neuron disease in adults, with the majority of individuals dying within 3-5 years of symptom onset. The disease is defined by the degeneration and death of motor neurons in the brain and spinal cord, resulting in progressive weakness and eventually death, typically from respiratory muscle weakness^3^. Around 5-10% of ALS cases are inherited, termed familial ALS (fALS), with the remaining cases considered sporadic ALS (sALS). After decades of study, the etiology of sALS remains a mystery, although suspected risk factors for ALS include exposure to heavy metals, pesticides, chemical solvents, cigarette smoke, and unidentified factors related to US military service^4–7^.

Along with these environmental risk factors, there has been a long history, with variable success, in the search for pathogens that might contribute to ALS^8–12^ and other neurodegenerative diseases such as Alzheimer’s disease (AD)^13–15^, Parkinson’s disease (PD)^16–18^, and multiple sclerosis (MS)^19^.

Diverse pathogens have been reported in the blood, cerebrospinal fluid (CSF) and central nervous system (CNS) from ALS patients. For example, bacteria that have been detected include *Cutibacterium acnes, Corynebacterium sp, Fusobacterium nucleatum, Lawsonella clevelandesis*, and *Streptococcus thermophilus* in CSF^20^, and mycoplasma in blood^21^. Fungi, including *Candida famata, Candida albicans, Candida parapsilosis, Candida glabrata*, and *Penicillium notatum*, have been detected in CSF, while *Malassezia globosa, Cryptococcus neoformans^11^*, and *Candida albicans* have been found in various regions of the CNS^11,22,23^. The search for viruses that contribute to ALS pathology is much more extensive and includes studies on herpes virus^9,24^, enterovirus^9,25-28^, human immunodeficiency virus (HIV)^29,30^, and human endogenous retrovirus (HERV-K)^31–33^. Importantly, multiple studies using immunohistochemistry have shown an increased load of various pathogens in ALS samples compared to controls in multiple tissues suggesting these pathogens are present and cannot be simply attributed to contamination^9,11,20,22,23^. Ultimately, the presence of ALS dysbiosis is unresolved and remains an active area of investigation with evidence for^34–38^ and against^39^ it.

The biological role that these alternative microbiotas play in ALS is also unclear. ALS patients may have a compromised blood brain barrier (BBB) or blood spinal cord barrier (BSCB) function^40,41^. It has been reported that ALS patients also have elevated Gram negative endotoxin/lipopolysaccharide (LPS) in the blood^42^. Patients with ALS also display activation of the innate immune system along with changes in blood^43,44^, spinal cord and motor neurons^45^, but if and how bacteria might influence activation is an active area of research. A “dual hit” hypothesis by Correia et al. suggests inflammation via LPS may contribute to mis-localization and aggregation of ALS-implicated protein TAR DNA-binding protein 43 (TDP-43)^46^.

Numerous studies have looked for biomarkers of ALS^47^ using metabolomics^48,49^, neuroinflammation^50,51^, DNA methylation^52,53^, gene expression^54^, microRNA expression^55,56^ and our previous study which analyzed protein levels of poly(GP) in *C9ORF72-associated* ALS (c9ALS)^57^. The search for pathogens using sequencing data from blood samples in ALS patients has been conducted before^58–61^, but previous efforts have not looked for novel pathogens.

Next-generation sequencing (NGS) technologies have shown broad detection of pathogens in a target-independent unbiased fashion^62–65^. However, designing a microbial detection experiment is non-trivial considering the variety of methods^66^ and algorithms^67^ that can be applied. Our primary goal when designing a new pipeline was to be conservative and unbiased with regards to discovery of novel pathogens. Furthermore, we wanted our pipeline to allow for the quantification of both novel and known pathogens. While other pipelines have used reads that do not map to the host genome (unmapped reads) for microbial identification and quantification, these pipelines cannot be used for discovery as they rely on existing databases of microbial genomes^68–71^. Thus, we opted for de-novo assembly of unmapped reads into contigs, followed by alignment of unmapped reads back to these contigs for quantification. A similar pipeline known as IMSA^72^ uses this strategy, but disregards contigs that might be identified by translated amino acid sequence similarity using BLASTX (a set we call the “dark biome”) as well as subsequent contigs with no BLASTN or BLASTX hit (a set we call the “double dark biome”).

We validated our pipeline by using datasets with known bacterial or viral infections. We also examined the differences in microbial identification between polyA and total RNA recovery in multiple tissues, and investigated the effects of globin depletion of blood samples. We then used our pipeline on a novel blood dataset (termed “Our Study”) as well as on five other published ALS datasets from blood or spinal cord samples, analyzed each dataset individually, and analyzed across datasets for changes in microbiota. While we did not identify any microbes enriched in the blood of ALS patients, we did identify and validate a novel virus-like sequence, demonstrating the potential of the bioinformatic pipeline we have established.

## MATERIALS AND METHODS

### Blood Collection and RNA Extraction

A total of 120 RNA whole blood samples constitute Our Study, which included 30 healthy controls (from general population that do not have blood relatives suffering from ALS, CTL), 30 pre-symptomatic *C9ORF72* mutant carriers (C9A), 30 symptomatic *C9ORF72* ALS cases (C9S), and 30 symptomatic *C9ORF72*-negative ALS cases (SYM). PAXgene blood RNA tubes were collected at Mayo Clinic Jacksonville and at University of Miami. All 120 RNA samples selected for RNA-seq were received and processed at Mayo Clinic Jacksonville using PAXgene blood RNA kit following manufacturer’s recommendations (Qiagen). Blood RNA was of high quality, assessed in an Agilent Bioanalyzer (Agilent), with RNA integrity values ranging from 7.4 to 9.8, with a median value of 8.7. RNA samples were then sent to The Jackson Laboratory for globin depletion, library preparation and sequencing of total blood RNA.

### Globin Depletion

Due to the abundance of large haemoglobin RNA transcripts present in the blood, a globin depletion step, using the Ambion GLOBINclear kit (AM1980), was performed before sequencing of the blood RNA samples in order maximize coverage on non-globin genes. In brief, one microgram of total RNA was used as starting material, and specific biotinylated oligos were used to capture globin mRNA transcripts. The capture oligos were hybridized with total RNA samples at 50°C for 30 min. Streptavidin magnetic beads were then used to bind to the biotinylated capture oligos hybridized to globin mRNA by incubating at 50°C for 30 min. The magnetic streptavidin beads-biotin complex were then captured to the side of the tubes by a magnet, and the resulting supernatant is free of globin mRNA. The globin depleted RNA was further purified by RNA binding beads and finally eluted in elution buffer. The resulting RNA free of >95% globin mRNA transcripts was then processed for next generation sequencing. Of note, to assess the efficiency of the globin RNA depletion, 10% of the samples processed were selected randomly and amplified using a Target-Amp Nano labeling kit (Epicentre). Samples were normalized to 100 ng input and reverse transcribed. First strand cDNA was generated by incubating at 50°C for 30 min with first strand premix and Superscript III. This was followed by second strand cDNA synthesis through DNA polymerase by incubating at 65°C for 10 min and at 80°C for 3 min. In-vitro transcription was then performed at 42°C for 4 hours followed by purification using RNeasy mini kit (Qiagen).

Due to the large number of samples, the globin depletion step was performed in two batches. We provided guidelines on how samples would be divided among the batches and also for how samples would be grouped in the sequencing runs in order to minimize technical variability. The Jackson Laboratory personnel were blinded to the identity of the samples.

RNA-seq of total blood RNA (globin and ribosomal RNA depleted) was performed in an Illumina HiSeq4000 with >70 million read pairs per sample. Raw reads were then sent back to us for bioinformatics analyses.

### Quantitative RT-PCR for blood RNA samples

A total of 500 ng of total blood RNA was used for reverse transcription polymerase chain reaction (RT-PCR), using the High Capacity cDNA Transcription Kit with random primers (Applied Biosystems). Quantitative real-time PCR (qRT-PCR) was performed using SYBR GreenER qPCR SuperMix (Invitrogen). Samples were run in triplicate, and qRT-PCRs were run on a QuantStudio 7 Flex Real-Time system (Applied Biosystems).

List of primers and their sequences:

*RDRP* forward 5’-GCTGTCAAATCGGTTTCCAAC-3’; *RDRP* reverse 5’-CTGCCTTCGTCATCTTGGAG-3’;

*GAPDH* forward 5’-GTTCGACAGTCAGCCGCATC-3’;

*GAPDH* reverse 5’-GGAATTTGCCATGGGTGGA-3’.

### Transcriptomics

See pipeline description in results for an overview of the pipeline; see bioinformatics supplement File S1 for a more detailed description of the analysis pipeline, versions, and statistical quantification. All data in this study was processed identically using the pipeline.

### Statistical Analysis

To assess statistical differences between conditions, a two tailed Student’s *t*-test is calculated using normalized coverage for contigs or binned normalized coverage for species/genus, etc. The number of contigs or genus/species is used to obtain an adjusted p-value using scipy in Python. Cutoff for statistical significance is less than an adjusted p-value of 0.05 unless otherwise stated.

### Data availability

Raw sequencing data for Our Study dataset is available in the NCBI Sequence Read Archive under the accession number (PRJN). All other datasets are publicly available and all of the code used in this manuscript is available at https://github.com/Senorelegans/MysteryMiner. Supplemental material available at figshare: https://doi.org/(INSERT).

## RESULTS

### Pipeline description

Mystery Miner is written as a Nextflow pipeline. Below is a short overview of the Mystery Miner pipeline (Fig1). A more in-depth explanation, list of software and versions used, and typical parameters of each step is described in the bioinformatics supplement, and all of the code used in this manuscript can be found at https://github.com/Senorelegans/MysteryMiner.

**Figure 1.**
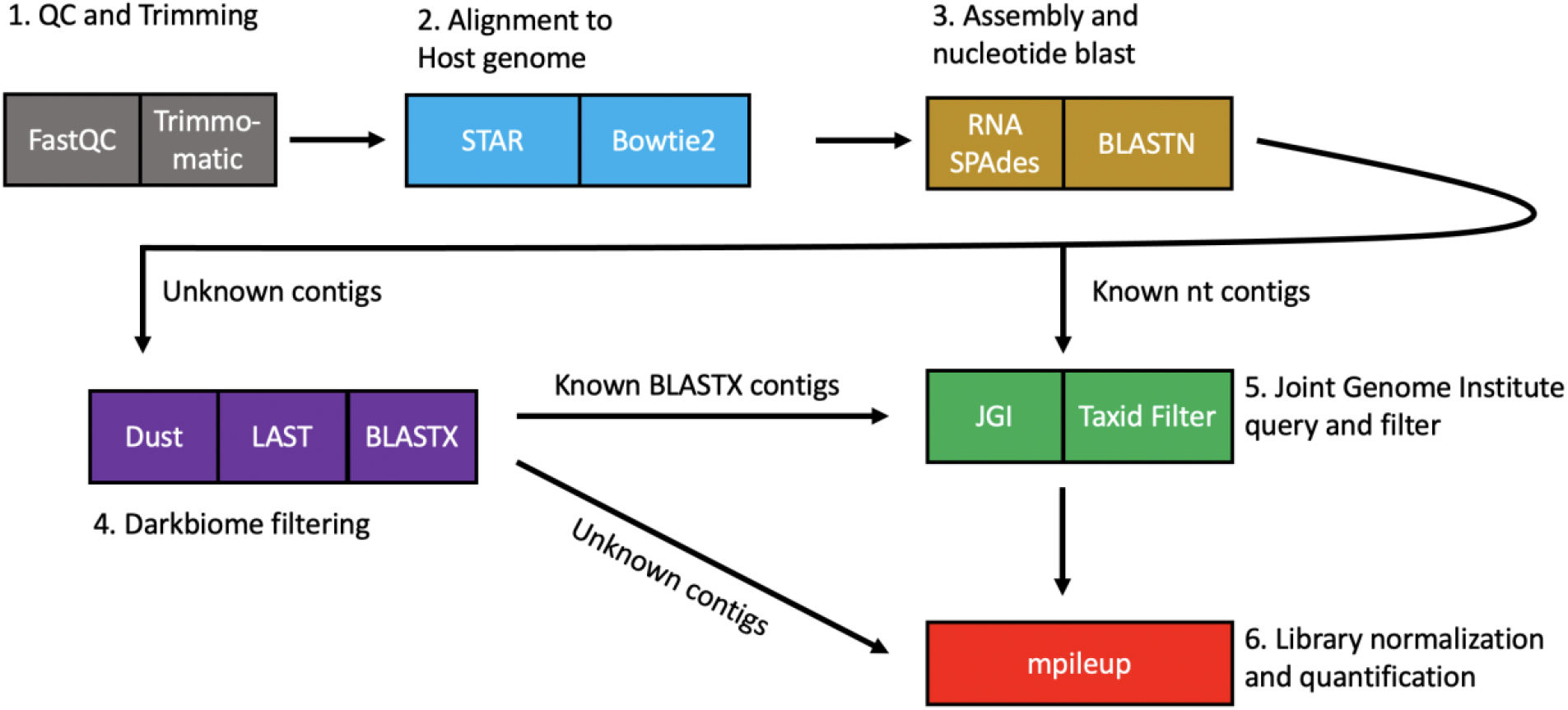
Diagram of Mystery Miner Pipeline. Reads were first checked with FastQC and trimmed using Trimmomatic (1. grey). Reads were then aligned to the host genome using various aligners (2. blue). Non-host (unmapped) reads were assembled into contigs with RNA SPAdes and regular biome contigs were identified with BLASTN (3. yellow). Unidentified contigs were filtered for repetitive sequences with Dust, filter by single, group or all with LAST, and dark biome contigs were identified with BLASTX. Double dark biome unidentified BLASTX contigs were sent directly to quantification (4. purple). Dark biome and regular biome contigs were assigned complete taxonomy using the JGI server and filtered one last time to remove mammalian/host genome contigs (5. Green). Non-host reads were then mapped to all contigs and normalized coverage was calculated for subsequent statistical analysis.

Raw reads were first checked for quality using FastQC then trimmed to remove both adaptor contamination and low quality basecalls using Trimmomatic. Trimmed reads were then mapped to the host genome using multiple alignment algorithms in series (STAR, Bowtie2) and unmapped reads were retained for contig assembly. Filtering out host reads made downstream assembly faster and required less memory. We assembled contigs from unmapped reads with the SPAdes assembler (with “-rna” setting). This assembler was chosen for its recent use in studies of microbial diversity^73^ and proven robustness to biological and technical variation^74^. The species each contig belongs to was identified with BLASTN using default settings, and the top hit for each contig was retained (a set we call “regular biome”). Contigs with no BLASTN hits were then filtered to remove highly repetitive regions (DUST) and retained if they had a greater than 60% pairwise alignment (LAST) between contigs assembled from a single sample, group/condition, or all samples.

We then identified contigs that lacked detectable nucleotide similarity to any GenBank entry but showed similarity at the amino acid level using BLASTX (“dark biome”). Furthermore, contigs with no BLASTN or BLASTX hits were labelled as “double dark biome” (also filtered by DUST and LAST). Every contig in the “regular biome” and “dark biome” were then queried against the Joint Genome Institute Server for additional taxonomic information. As Mystery Miner is an agnostic tool, it cannot distinguish between true tissue or cell-associated microbes and experimentally introduced contamination.

For quantification, we mapped the non-host reads using Bowtie2 to the contigs obtained from SPAdes. Next, we mapped reads to contigs using samtools mpileup (default mapq score) to calculate the amount of reads over each base pair in a contig. We then calculated coverage on the contigs by summing all of the counts for each base pair in a contig and dividing by the length of the contig. We then calculated normalized coverage by library size using the number of mapped reads to the host genome. This gave us normalized coverage (NC) for a contig or binned normalized coverage (BNC) for multiple contigs within a species/genus, etc. To assess statistical differences between conditions, a Student’s *t*-test was calculated through NC or BNC, using the number of contigs or genus/species to obtain an adjusted p-value using scipy in Python.

### Validating Mystery Miner on datasets with known bacterial or viral infection

To confirm that Mystery Miner is able to recover and quantify known bacterial infections from sequencing data, we utilized an *in vitro* model of *Chlamydia trachomatis* infection (Humphrys 2016)^75^. In this study, epithelial cell monolayers were infected with *Chlamydia trachomatis*; and polyA RNA (6 samples) and total RNA (6 samples) were sequenced 1 hour and 24 hours post infection (hpi). Using the Mystery Miner pipeline, out of 5.32 × 10^6^ reads from all of the samples, 6.04 × 10^5^ reads remained unmapped (~11.34%) after trimming and mapping to the host genome (File S2). From these non-host reads, 3,257 contigs were assembled and 1,199 of these contigs were identified as regular biome (File S3). An additional 27 contigs had no BLASTN hit. Of these, we identified 2 dark biome (BLASTX identified) and no double dark biome (no BLASTX or BLASTN hit) contigs (File S4 and File S5).

Using Mystery Miner we successfully identified, and found significantly elevated levels, of *Chlamydia trachomatis* (BNC by species) in 24 hours post infection (hpi) samples compared to 1 hpi samples in both polyA (Padj = 0.004) and total RNA (Padj = 0.0005). In addition to *Chlamydia trachomatis*, we identified 6 additional bacterial species and one viral species (Alphapapillomavirus 7) in the samples (Figure 2A), including significantly elevated levels of *Mycoplasma hyorhinis* contigs in total RNA samples. No significant differences were observed in the dark or double dark contigs (File S6).

**Figure 2.**
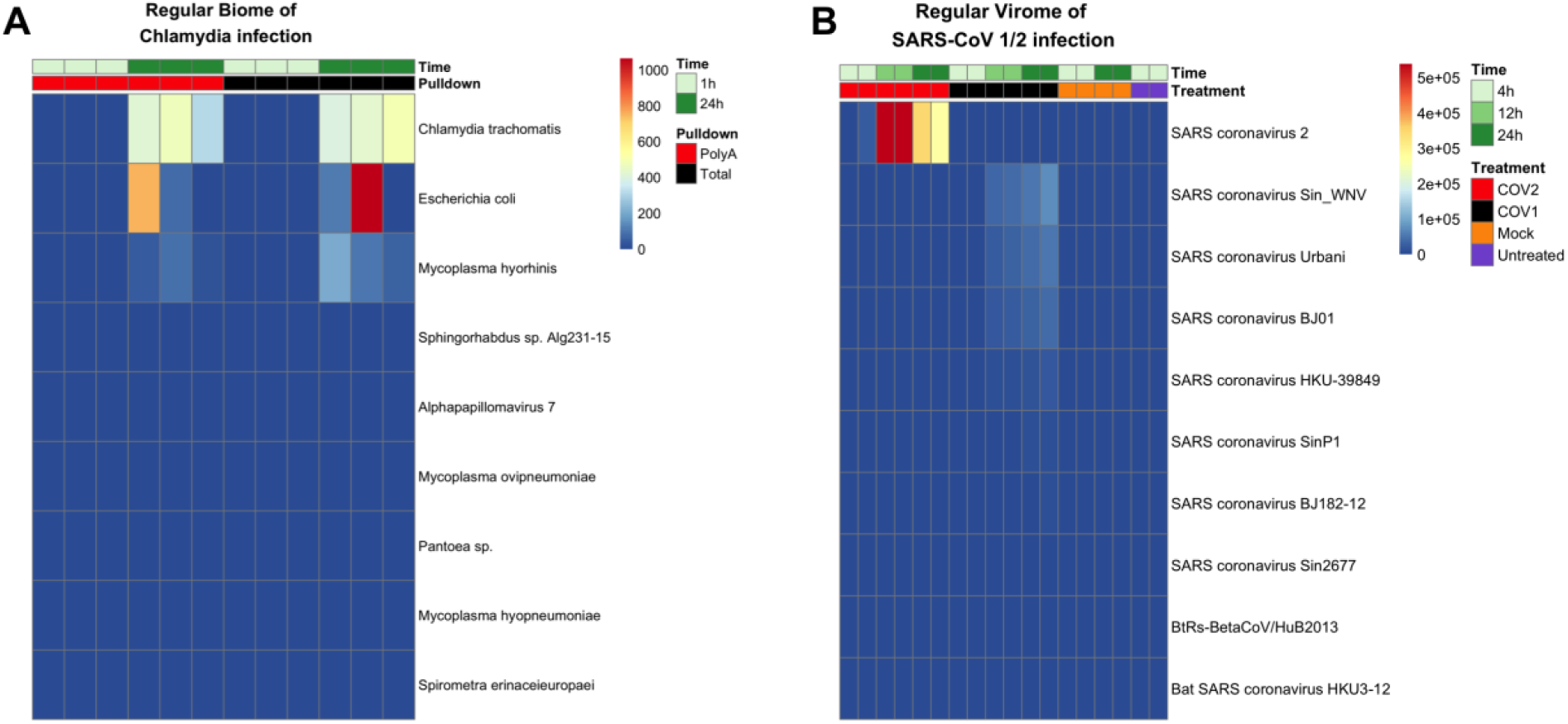
Heatmap of binned normalized coverage for bacterial or viral infected datasets. **A.** Regular biome contigs binned by species from Humphrys et al., 2016. Time refers to 1or 24 hours post infection (hpi) of epithelial cell monolayers with *Chlamydia trachomatis* (green). Pulldown refers to library enrichment for polyA RNA (red) or total RNA (black). **B**. Regular virome of contigs binned by name from Emanuel et al., 2020 for SARS-CoV-2 infected cells (COV2) (red), or SARS-CoV-1infected cells (COV1) (black), mock virus (orange), or untreated sample (purple). Time refers 4,12, or 24 hpi of Calu3 cells with indicated virus (green). Top 10 hits per experiment shown for brevity.

To confirm that the pipeline can detect known viral infections, we ran Mystery Miner on a total RNA dataset from an *in vitro* model of severe acute respiratory syndrome coronavirus (SARS-CoV) 1 or 2 infection (Emanuel2020^76^). In this study human epithelial Calu3 cells were infected with SARS-CoV-1 or SARS-CoV-2 (4, 12, or 24 hours), mock (4 or 24 hours), or untreated (4 hours).

Out of the 2.81 × 10^8^ reads obtained from all of the samples, 8.23 × 10^7^ reads remained unmapped (~29%) after trimming and mapping to the host genome (File S2). From these non-host reads, 42,816 contigs were assembled, of which 1,346 regular biome, 27 dark biome, and 7 double dark biome contigs passed the filtering steps (File S2, File S3, File S4, File S5)

Mystery Miner successfully identified both SARS-CoV-2 and SARS-CoV-1 isolates and found significantly elevated levels of each virus compared to controls (Figure 2B). Hereafter we refer to SARS-CoV-1 or SARS-CoV-2 infected cells as COV1 or COV2 to avoid confusion with recovered names of contigs. Consistent with the viruses being similar, we identified both SARS-CoV-2 and SARS-CoV-1 in both the COV1-24hr and COV2-24hr samples when compared to mock-24hr. However, when we compared COV2-24hr to COV1-24hr, we were able to distinguish SARS-CoV-1 isolates from SARS-CoV-2 in the appropriate samples (i.e., SARS-CoV-2 was significantly elevated in COV2). Similar results were seen in the 12 hour samples but the 4 hour samples did not have sufficient viral reads to detect either SARS-CoV virus (File S7). To simulate a novel virus, we ran Mystery Miner on versions of the BLASTN and BLASTX databases obtained before SARS-CoV-2 was discovered and were able to properly identify SARS-CoV-2 as a bat related coronavirus^77^ (Figure S1) (File S7).

Collectively, these data show that Mystery Miner is able to identify potential bacterial and viral infections, properly identify infected samples using quantification, and detect significant differences between infected samples and controls for bacteria, viruses, and isolates of a virus.

### Effects of library pulldown or globin depletion in RNA-seq datasets

In order to compare effects of library enrichment or depletion, we compared recovered pathogens in a dataset that has polyA enrichment or rRNA depleted total RNA from blood or colonic tissue (VonSchack2018)^78^. When we compared polyA RNA vs total RNA and looked at BNC by superkingdom of bacteria we found significantly more reads map to bacteria for total RNA than polyA RNA (Padj = 0.0349), in blood but not in colon (Padj=0.11709) (Figures S2 and File S8). We found similar amounts of significant BNC by species for polyA RNA vs total RNA in blood (29) and in colon (26). We then looked at significant BNC by genus and found double the amount in blood (14) compared to colon (7), with only one significant genus (*Actinomyces*) found in both comparisons. We did not find any significant differences in coverage when we looked at the species, genus or superkingdom level for viruses (File S8). We conclude that library enrichment with total RNA compared to polyA RNA increases bacterial recovery and diversity in blood.

We next looked at a RNA-seq dataset from whole blood with globin depleted (GD) vs non-globin depleted (NGD) total RNA (Shin2014^79^). With BNC by superkingdom, we found significantly increased levels in globin depleted vs. not-depleted samples for both bacteria (Padj = 0.004) (Figure S3) and viruses (Padj = 0.030) (Figure S4). We also found significant differences in BNC by species (Figure S5) or genus (Figure S6) primarily from *E. coli* with elevated levels in globin-depleted blood RNA. We did not find any significant differences when we looked for viruses at the species or genus level (File S9).

### Analysis of Our Study

We used Mystery Miner on our novel RNA-seq dataset of globin depleted and rRNA depleted total blood RNA from 120 individuals. These samples were from four subject groups including healthy control participants (CTL), ALS symptomatic *C9ORF72* negative patients (SYM), *C9ORF72* positive ALS symptomatic patients (C9S) and *C9ORF72* positive asymptomatic individuals (C9A).

The entire dataset contains a combined 8.64 × 10^9^ reads. Approximately 2.7% (2.34 × 10^8^) of the reads did not map to the human genome. From these non-host reads 2,976,988 contigs were assembled and 17,047 BLASTN contigs (regular biome) were identified. A total of 25,815 contigs had no BLASTN hit and after filtering we identified 2,980 dark biome (BLASTX identified) and 859 double dark biome (no BLASTX or BLASTN hit) contigs (File S2, File S3, File S4, File S5).

In general, we found a modest positive correlation between library size and number of bacterial contigs assembled, species detected (Figure 3), and genera detected for each sample as well as a homogenous mixture of samples with respect to disease status. No specific taxonomy or contig sequence correlated with participant class within the dataset. Yet, by pooling bacterial read counts across all of the samples, we found *alpha proteo-bacteria, Actinobacteria, Firmicutes*, and *Bacteroidetes* as the most highly represented taxonomies, consistent with other blood biome studies^80^ (Figure S6). Most of the bacterial genera (~65%) assigned to the dark biome contigs were also found in the regular biome, however this was not the case in the viral sets, as only 5% (4/78) of dark viral contigs were observed in the regular biome (File S10). This observation suggested that our pipeline might be identifying novel viral sequences.

**Figure 3.**
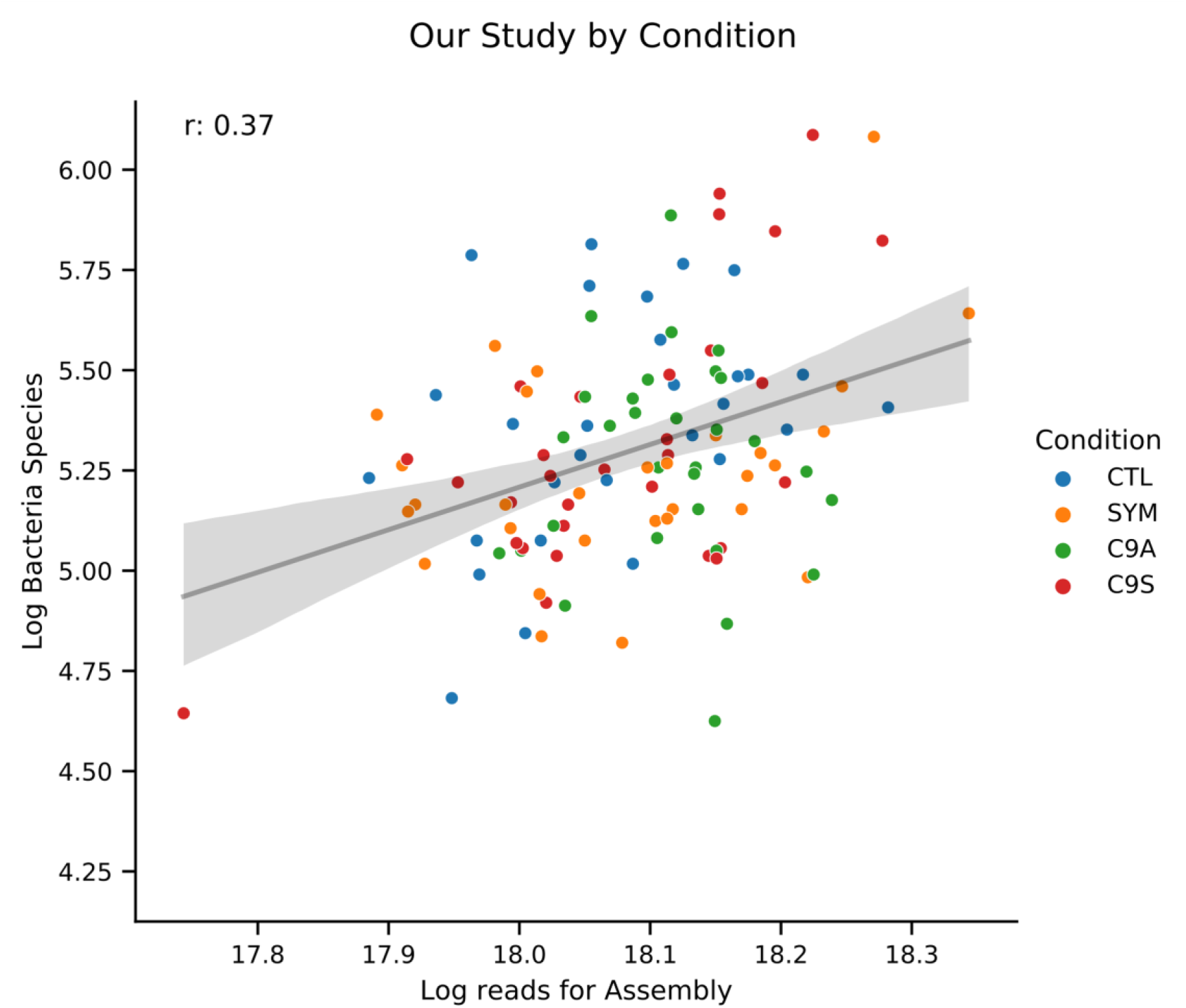
Log number of bacterial species vs Log reads for Assembly in Our Study. Scatterplot where each dot is a sample from a dataset with log number of bacterial contigs assembled on the Y-axis and Log reads used in SPAdes on the X-axis. Samples show a modest correlation (Pearson’s r=0.37) between library size and bacterial species recovered. Data fit with a regression (black line) and 95% confidence interval (gray area).

Within the dark biome contigs, we noted numerous contigs with a region of protein sequence similarity to RNA-dependent RNA polymerase (RdRP) from multiple RNA viruses, showing highest similarity to the velvet tobacco mottle virus (first row in heatmap of Figure 4, complete metadata shown in Figure S7). Our attention was drawn to the largest (~5 kb) dark biome contig (one of the contigs showing similarity to the velvet tobacco mottle virus) hereafter labeled as “RDRP contig”. To confirm the presence of the RDRP contig in the original samples, we designed primers to the RDRP contig and performed reverse transcriptase polymerase chain reaction (RT-PCR) on seven samples, four of which had high coverage (predicted present) and three with zero coverage (predicted absent). We found typical levels for detection of a virus^81^ in the samples with high coverage and detected nothing in samples with zero coverage (Table 1). We conclude that Mystery Miner is biologically validated and can recover unknown pathogens from human subjects.

**Figure 4.**
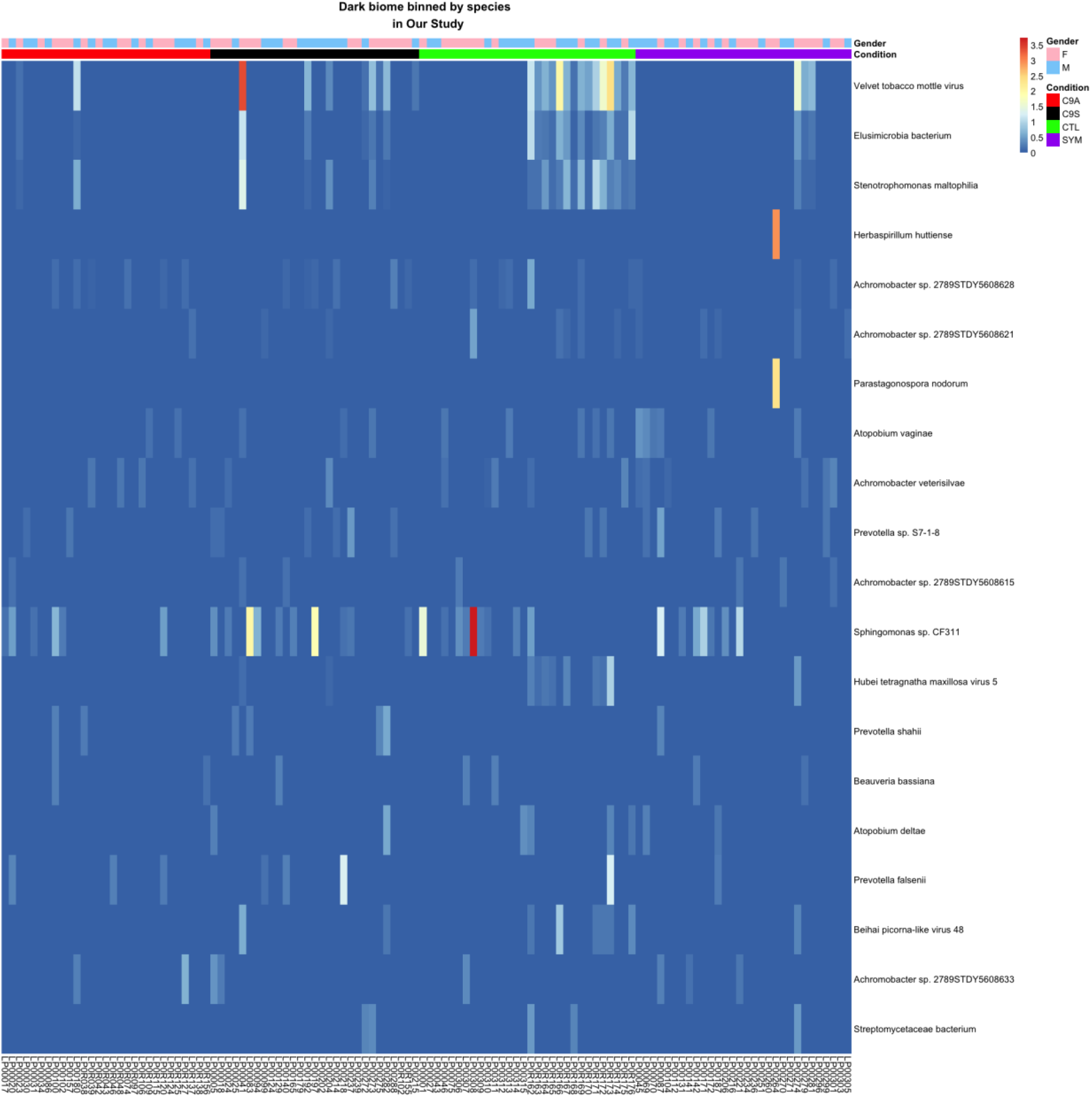
Heatmap of dark biome contigs binned by species in Our Study. Heatmap of normalized coverage of dark biome contigs binned by species. The highest coverage belongs to contigs that show high similarity to velvet tobacco mottle virus. Zero coverage is blue and goes to red with increasing values. These samples were from four subject groups including healthy controls [(CTL) green], *C9ORF72* negative ALS symptomatic [(SYM) purple], *C9ORF72* positive ALS symptomatic [(C9S) blue] and *C9ORF72* positive asymptomatic [(C9A) red] patients. Sex indicated as light blue (male) and pink (female). Top 20 species sorted by binned normalized coverage was shown for brevity.

**TABLE 1.** RT-PCR AND NORMALIZED COVERAGE RESULTS FOR RDRP CONTIG. Quantitative RT-PCR and normalized coverage results from the 5180 bp RDRP contig. For the RDRP contig positive samples (top 4) with high normalized coverage and detectable Ct values and negative samples (bottom 3) with no normalized coverage and undetectable Ct values. GAPDH was used as a positive control for qRT-PCR and shows comparable levels for all samples. These samples were from three conditions *C9ORF72* negative ALS symptomatic patients (SYM), *C9ORF72* positive ALS symptomatic patients (C9S) and *C9ORF72* positive asymptomatic individuals (C9A).

| Condition | Sample | GAPDH<br>RT-PCR<br>Ct Value | RDRP<br>RT-PCR Ct<br>Value | RDRP<br>RNA-seq<br>Normalized<br>Coverage |
| --- | --- | --- | --- | --- |
| SYM | LP00274 | 20.562019 | 36.401 | 1.56 |
| C9S | LP00041 | 20.783213 | 36.346 | 3.39 |
| C9S | LP00192 | 20.899612 | 35.636 | 0.67 |
| C9A | LP000180 | 19.982108 | 34.832 | 1.11 |
| C9S | LP000183 | 20.176418 | undetermined | 0 |
| C9S | LP000197 | 20.125161 | undetermined | 0 |
| C9A | LP000157 | 20.062433 | undetermined | 0 |

### Analysis of published ALS datasets

We next sought to explore whether similar results would be obtained from other ALS datasets. To this end, we examined five other publicly available ALS datasets, consisting of two that used total RNA from blood (Linsley2014^82^, Gagliardi 2018^58^), and three datasets from spinal cord (Brohawn2016^83^, Ladd2017^84^, Brohawn 2019^85^). We have provided a summary table of datasets for all studies used in this paper (Table 2). As we observed in Our Study, we first noted that increased library size correlated with an increased number of bacterial contigs assembled, species detected, and genera detected (Figure 5, and Figure S8-10 show all datasets used in this study).

**Figure 5.**
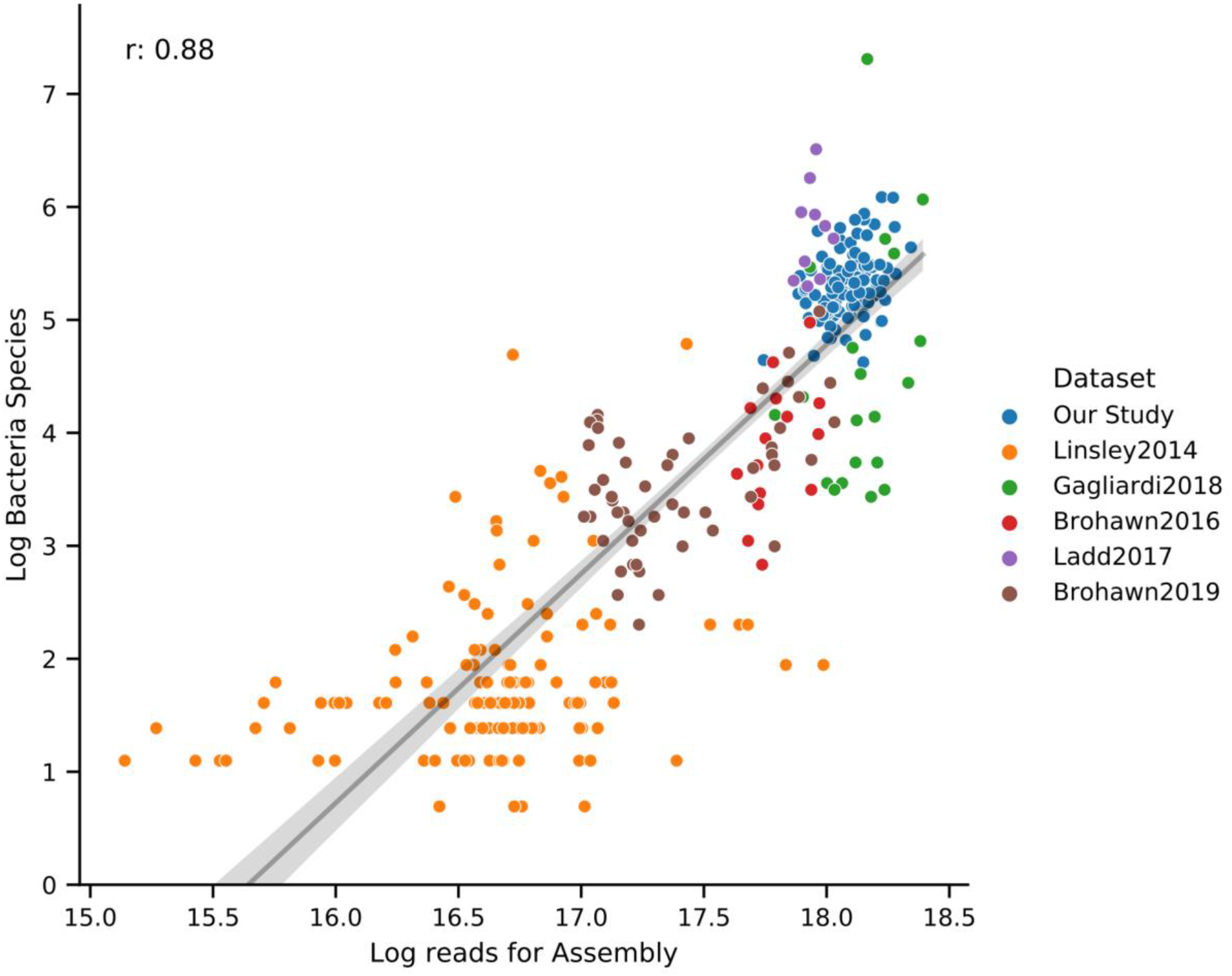
Log number of bacterial species vs Log reads for Assembly for ALS Datasets. Scatterplot where each dot is a sample from a dataset with log number of bacterial contigs assembled on the Y-axis and Log reads used in SPAdes on the X-axis. ALS datasets show a high correlation (Pearson’s r=0.88) between library size and bacterial species recovered. Data fit with a regression (black line) and 95% confidence interval (gray area).

**TABLE 2.** STUDY DESIGN FOR DATASETS USED IN THIS PAPER. Overview of the datasets used in this paper. The first three studies are only used to validate our pipeline. The six subsequent studies are ALS related from both blood and spinal cord.

| Name | Groups | # Samples | Tissue | Pulldown |
| --- | --- | --- | --- | --- |
| Humphrys2016 | 1- or 24-hours post infection with *Chlamydia trachomatis* | 12 | Cultured epithelial cell monolayers | PolyA<br>Total RNA |
| VonSchack2018 | PolyA or Total RNA from blood or colon | 16 | Whole Blood<br>Colon | PolyA RNA<br>Total RNA |
| Shin2014 | Globin depleted<br>Not globin depleted | 24 | Whole Blood | Total RNA |
| Emanuel2020 | Severe acute respiratory syndrome coronavirus 1 or 2 infection<br>Controls | 18 | Calu3 cells | Total RNA |
| Our Study | *C9ORF72* negative ALS,<br>*C9ORF72* positive and symptomatic ALS,<br>*C9ORF72* positive asymptomatic participants<br>Controls | 120 | Whole Blood | Total RNA<br>hemoglobin and rRNA depleted |
| Linsley2014 | ALS<br>type 1 diabetes, sepsis, multiple sclerosis patients before and 24 hours after the first treatment with IFN-beta<br>Controls | 134 | Whole blood | Total RNA |
| Gagliardi2018 | Sporadic ALS, ALS with mutations in *FUS*, *SOD1*, *TARDBP*<br>Controls | 20 | Peripheral blood mononuclear cells | Total RNA |
| Brohawn2016 | ALS<br>Controls | 15 | Cervical spinal cord | Total RNA<br>rRNA depleted |
| **Ladd2017** | **ALS**<br>**Controls** | **10** | **Laser capture**<br>**microdissection**<br>**(LCM) to isolate**<br>**cervical spinal cord**<br>**motor neurons** | **Total RNA** |
| **Brohawn2019** | **ALS, Alzheimer's**<br>**disease (AD),**<br>**Parkinson's disease**<br>**(PD)**<br>**Controls** | **53** | **Cervical spinal**<br>**cord** | **Total RNA** |

We then looked at the total overlap of genus or species to determine if there are similarities in recovered microbial or viral sequences between datasets. For genus in the regular bacteriome, our dataset had the highest number of unique genus (185), followed by Ladd2017 (117), and Gagliardi2018 (38). The highest number of overlapping bacterial genus was between our dataset and Ladd2017 (133) followed by the intersection between our dataset, Ladd2017 and Gagliardi2018 (61) and there was a modest overlap (24) for 9/10 datasets (Figure 6). We observed roughly the same trend in the regular bacterial biome at the species level and in the dark bacterial biome (S Figure 11, File S11). In contrast, the regular virome of each dataset was relatively unique with very low amounts of overlap (<= 3) between datasets (species and genus shows a similar pattern). Interestingly, the highest overlap for species in the dark virome was between our dataset and Ladd2017 (13), one of which is similar to RDRP viruses, although the contigs in Ladd’s data were not similar to the velvet tobacco mottle virus in our dataset (Figure S12, File S12).

**Figure 6.**
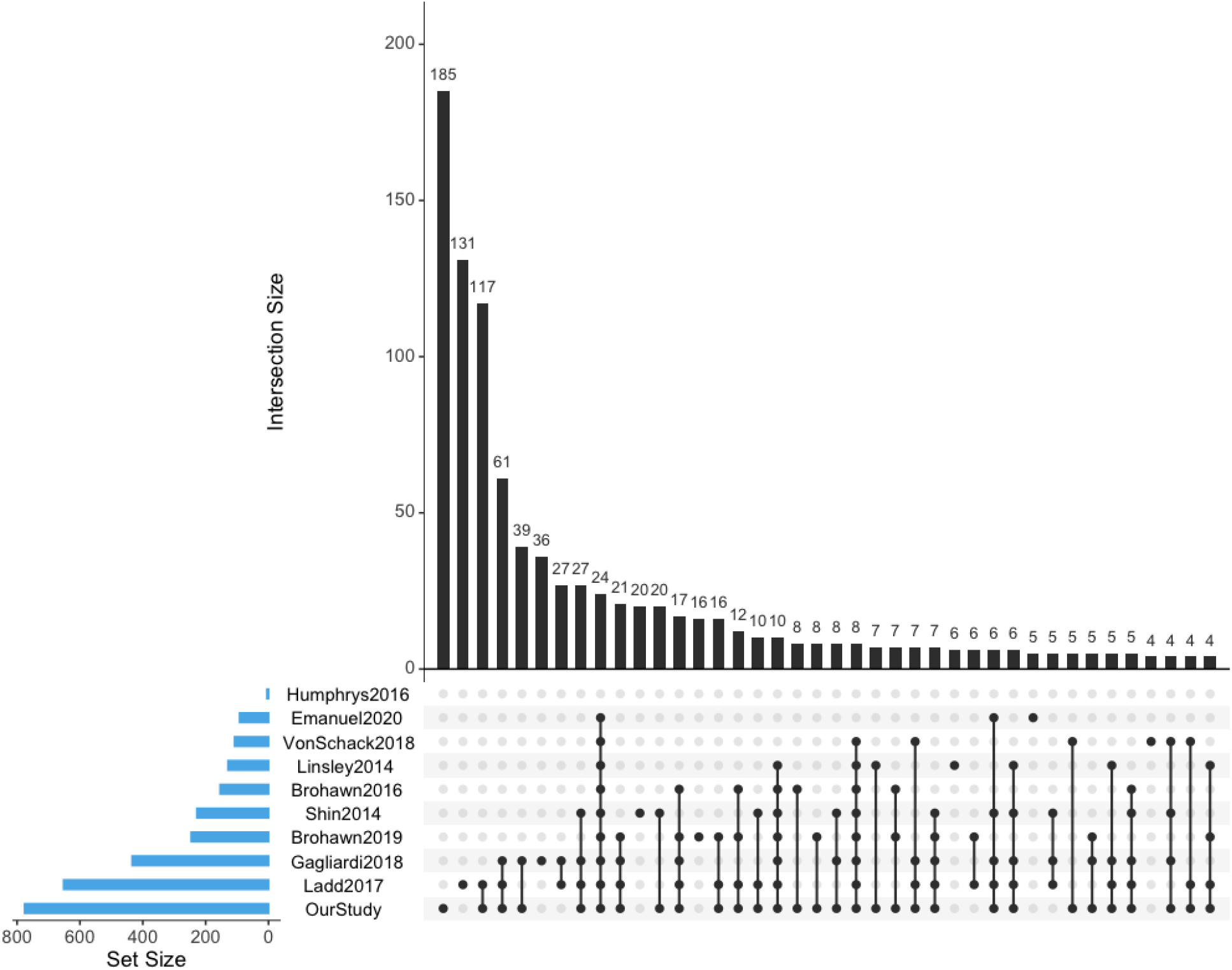
Upset plots of overlapping genus in the regular bacteriome between datasets. Upset plots are Venn diagram-like plots. A set refers to a dataset used in this study and each set is on a row with total amounts in a set as a blue bar plot on the left (ordered by set size). The black histogram on top shows the counts that are in the intersection of sets (a single dot for one dataset or connected dots for overlap of multiple datasets). Intersections less than 4 are removed for visualization purposes.

In addition to comparing datasets using taxonomy, we also compared contigs between datasets for nucleotide similarity (> 70%) using LAST (File S1 for methods). We found that in general, datasets in the regular biome with the largest amount of contigs have the most overlap. Unsurprisingly, in the dark biome we observed less overlap by nucleotide similarity and that our RDRP contig does not share nucleotide similarity with contigs from any dataset. In addition, we also compared the nucleotide similarity of double dark biome contigs and found there is not a large percentage of similar contigs between datasets (File S13).

### Comparison of taxonomy by condition within ALS datasets

Finally, we looked for differences in ALS vs control samples for each dataset. In the Gagliardi2014 dataset, when we compared ALS patients with the *FUS* mutation to controls, we found 3 significant differences in BNC by species in the regular bacteriome (*Neisseria sp., Pseudomonas sp., Sphingomonas sp.*) and one significant difference in BNC by genus in the dark bacteriome (*photobacterium*). In ALS patients with mutations in *SOD1* compared to controls, we found two species significantly different in the regular bacteriome (*Hydrogenophaga crassostreae, Sphingomonas hengshuiensis) (Gagliardi FUS and SOD1 supplement*). We did not find anything significant in sporadic ALS, or in ALS patients with *TARDBP* mutations with regards to genus/species (regular or dark biome or viruses) for Gagliardi2014. We found no significant statistical differences between ALS and control samples for genus/species of viruses/bacteria in the regular/dark biome for any of the remaining ALS datasets.

### Meta analysis between datasets

Since our dataset and many others had no significant comparisons for ALS vs control groups within each dataset, a meta-analysis between datasets using this criteria would be difficult. As a second pass analysis we created a less stringent filtering method in order to compare the presence of microbes for each group between datasets (ALS vs. ALS; or controls vs. controls) (Figure 7). We assigned a contig to a condition if ≥ 2 samples from that condition contain at least 90% of the summed normalized coverage (from all samples) to the contig. This filtering greatly reduced the number of comparable genus/species for each dataset and, for example, reduced the genus of the regular bacteriome in our dataset from 305 for all samples to 33 (SYM:8, C9S:6, C9A:2, CTL:17) (File S14).

**Fig 7.**
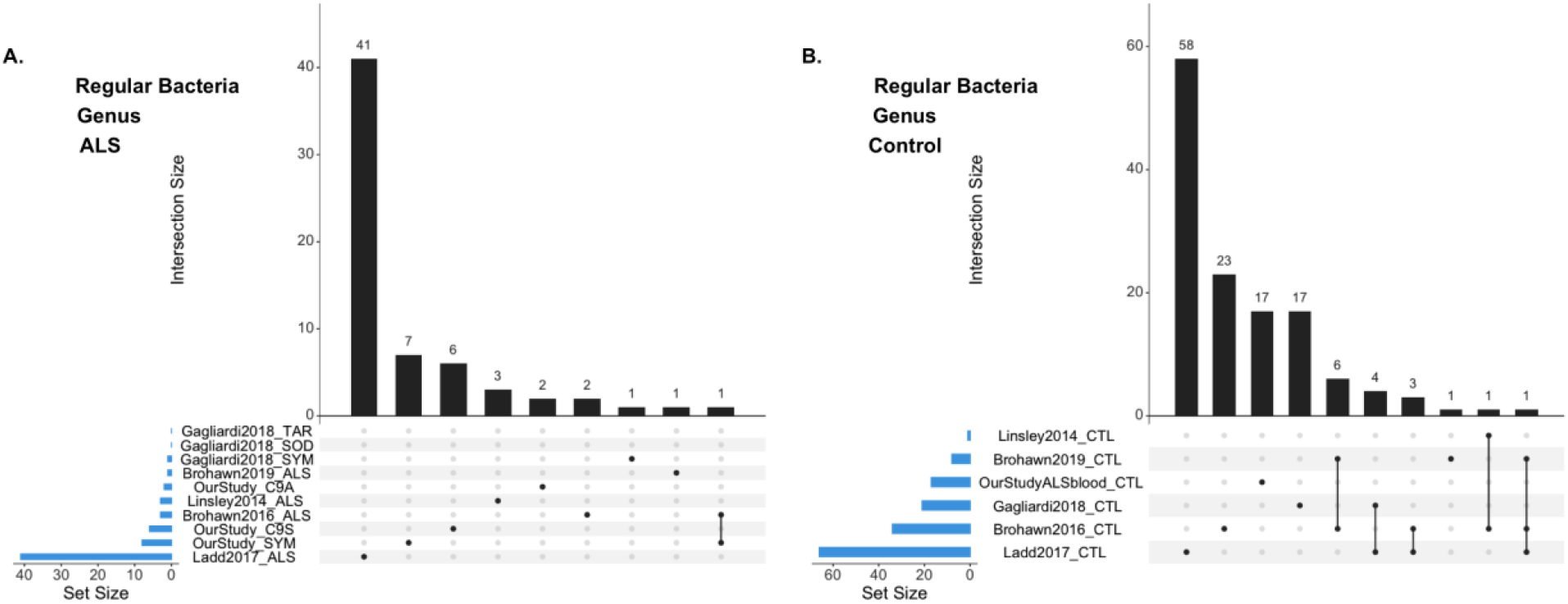
Upset plots of overlapping genus between datasets in the regular biome for ALS or controls. Upset plots are Venn diagram-like plots. A set refers to a contig that was assigned to a condition from a dataset. Each set is on a row with total amounts in a set as a blue bar plot on the left (ordered by set size). The black histogram on top shows the counts that are in the intersection of sets (a single dot for one dataset or connected dots for overlap of multiple datasets). A. ALS contigs in the regular bacteriome. B. Control contigs from the regular bacteriome.

When we looked at ALS or control contigs in the regular bacteriome, the highest number of unique genus or species was from Ladd2017, and in general there was a small amount of overlap between datasets (≤1 for ALS or ≤ 8 for controls) (Figure 7). When we looked at genus in the dark bacteriome we saw no overlap for ALS contigs and low overlap (≤ 1) between control conditions (species was similar) (File S14). In the regular virome there was no overlap between datasets and only our study (one contig from ALS) and Ladd2017 (three from ALS, five from controls) had contigs that passed the filter (similar values for species). When we looked in the dark virome by genus there was no overlap between datasets, and our dataset had only one genus (*Sobemovirus* from controls) with the rest coming from Ladd2017 (18 from controls, 5 from ALS) (File S15). In conclusion, despite our conservative and loose approaches, we did not find any convincing evidence in ALS samples that the presence (or lack of presence) of an organism (or multiple organisms) was different compared to control samples.

## Discussion

We have created Mystery Miner to search for and quantify known and unknown microbes in RNA-seq datasets as a tool for researchers to study dysbiosis and identify infectious agents. We validated the pipeline by recovering and quantifying *Chlamydia* and SARS-CoV reads from RNA-seq datasets from intentionally infected cells. Interestingly, we also identified *Mycoplasma* reads in the *Chlamydia* dataset, suggesting that Mystery Miner may also be able to identify unsuspected bacterial infections. We also use published data to investigate the difference of polyA vs total RNA recovery of bacterial species in multiple tissues. Perhaps surprisingly, we did not see a consistent difference in the recovery of bacterial reads between the two types of RNA-seq libraries, considering that bacterial transcripts are not expected to be polyadenylated. However, it is well-recognized that polyA selection is imperfect, and libraries constructed from polyA-selected RNA routinely contain transcripts thought not to be polyadenylated (e.g., rRNA). We also found increased recovery of bacterial species with globin RNA depletion in blood. This could be the result of an effective increase in read depth for bacteria when not sequencing globin, or an increase in contamination from the globin depletion step. We stress that our bioinformatic approach alone cannot distinguish between contamination and the true existence of microbial sequences in human tissue.

We then used Mystery Miner on Our Study dataset consisting of 8.64 × 10^9^ reads. This dataset was generated from whole blood total RNA that was depleted for both ribosomal and globin transcripts. It encompasses samples from controls, participants with a *C9ORF72* hexanucleotide expansion (symptomatic and pre-symptomatic), and *C9ORF72* negative ALS patients. We found no statistical difference in microbial sequence read coverage between controls and any class of ALS patients, examining either individual contigs or using various taxonomical binning approaches. We also did not detect any batch effects or obvious age-or sex-biases in the recovery of microbial reads (Figure S7). Overall, we found no evidence of blood microbial sequences contributing to either *C9ORF72* negative ALS or symptomatic patients harboring the *C9ORF72* hexanucleotide expansion. However, ALS is a CNS disease, so our findings in these blood samples do not necessarily preclude a role for microbes in this disease.

A unique aspect of Mystery Miner is that it tracks non-human reads that do not have significant BLASTN hits in GenBank. We were intrigued by the identification of a large contig (>5kb) in the dark biome of our ALS dataset that showed protein sequence similarity to RNA-dependent RNA polymerases, the essential replicase of RNA viruses. To validate that this virus-like sequence was not an artifact of contig assembly or a contaminant introduced during library construction or sequencing, we used RT-PCR of the original patient samples to demonstrate that this sequence was present in positive samples identified through the RNA-seq analysis and not detectable in negative samples. We cannot extrapolate from this specific example to determine what fraction of the “dark” and “double dark” sequences represent true novel microbial sequences present in human blood, but we note that analysis of human cell free blood DNA has reported the identification of thousands of novel bacterial sequences^86^. <u>We suggest that Mystery Miner is a generally useful tool for the identification of novel microbial sequences in RNA-seq data</u>.

To extend our analysis we processed publicly available blood and spinal cord ALS datasets through our pipeline. As observed in our dataset, library size generally correlated with number of bacterial contigs assembled and number of bacterial genera/species recovered. When the microbial sequences we found in our dataset were compared to the other datasets we found similar genera/species and, not surprisingly, larger datasets generally had greater overlap. One dataset (Ladd2017) yielded comparable recovery of bacteria and viruses for the regular biome but a far greater recovery bacteria and viruses in the dark biome compared to all the other datasets. This study used laser capture microdissection (LCM) to isolate cervical spinal cord motor neurons and had comparable read amounts per sample to other studies and was conducted in the same laboratory as two other studies (Brohawn2016, Brohawn2019). We are unsure why this dataset yielded a much larger dark biome compared to the other datasets. Potentially these differences are a byproduct of using LCM to acquire samples.

We then analyzed several publicly available ALS datasets for statistically significant differences between recovered microbial sequences in ALS and control samples. Only one dataset (Gagliardi2018) had any significant microbial sequence differences between control and ALS samples, specifically ALS patients with *FUS* or *SOD1* mutations. However, the sample number in this sub-study was small (N = 2-3), and in the case of the *SOD1* patients the excess microbial reads were in the control samples. In the absence of additional information (e.g., batch assignments for the samples) it is difficult to conclude that these sequence/sample correlations are meaningful. Finally, we compared identified microbial sequences in the control and ALS samples across the datasets and did not identify any genera/species that were preferentially recovered in either sample type.

Using our bioinformatic analysis pipeline Mystery Miner, we have not identified an association between ALS pathology and sequences corresponding to known or unknown microbial species. However, there are intrinsic limitations in using “repurposed” RNA-seq data to assay tissue-associated microbial sequences, including the relatively small number of non-human reads (<1% of total) upon which the analysis is based. This limited sequence signal could preclude identification of rarer microbes. Perhaps more problematic is the possibility that contaminating sequences obscure true tissue-associated microbial sequences. Any candidate microbes identified using Mystery Miner that correlate with human phenotypes will necessarily require independent validation. Despite these limitations, we believe Mystery Miner will be a useful tool for future researchers investigating known and unknown microbes that could contribute to disease, as our analyses have shown it to be sensitive to bacterial/viral agents in sequencing data.

## Acknowledgements

We thank the Jackson Laboratories for globin depletion and RNA-seq to generate Our Study dataset. This work was supported by the NIH (NS063964, to CL); the ALS Association (ALSA, to LP; #WA1096 and 16-IIP-253 to MP; and #2015 to MB); NINDS (R35NS097273, P01NS084974, to LP); the Mayo Clinic Foundation (to LP); Neuroscience Focused Research Team Mayo Clinic grant (to MP); the Association of Frontotemporal Dementia (AFTD, to LP); the Alzheimer’s Association-AD Strategic Fund (to LP); the Muscular Dystrophy Association (MDA #172123 to MB); the ALS Recovery Fund (to MB); Kimmelman Estate (to MB); and the Department of Defense (Chem-Bio Diagnostics program Grant HDTRA-1-18-1-0032 to RDD).

## Supplemental Figures

**Figure S1.**
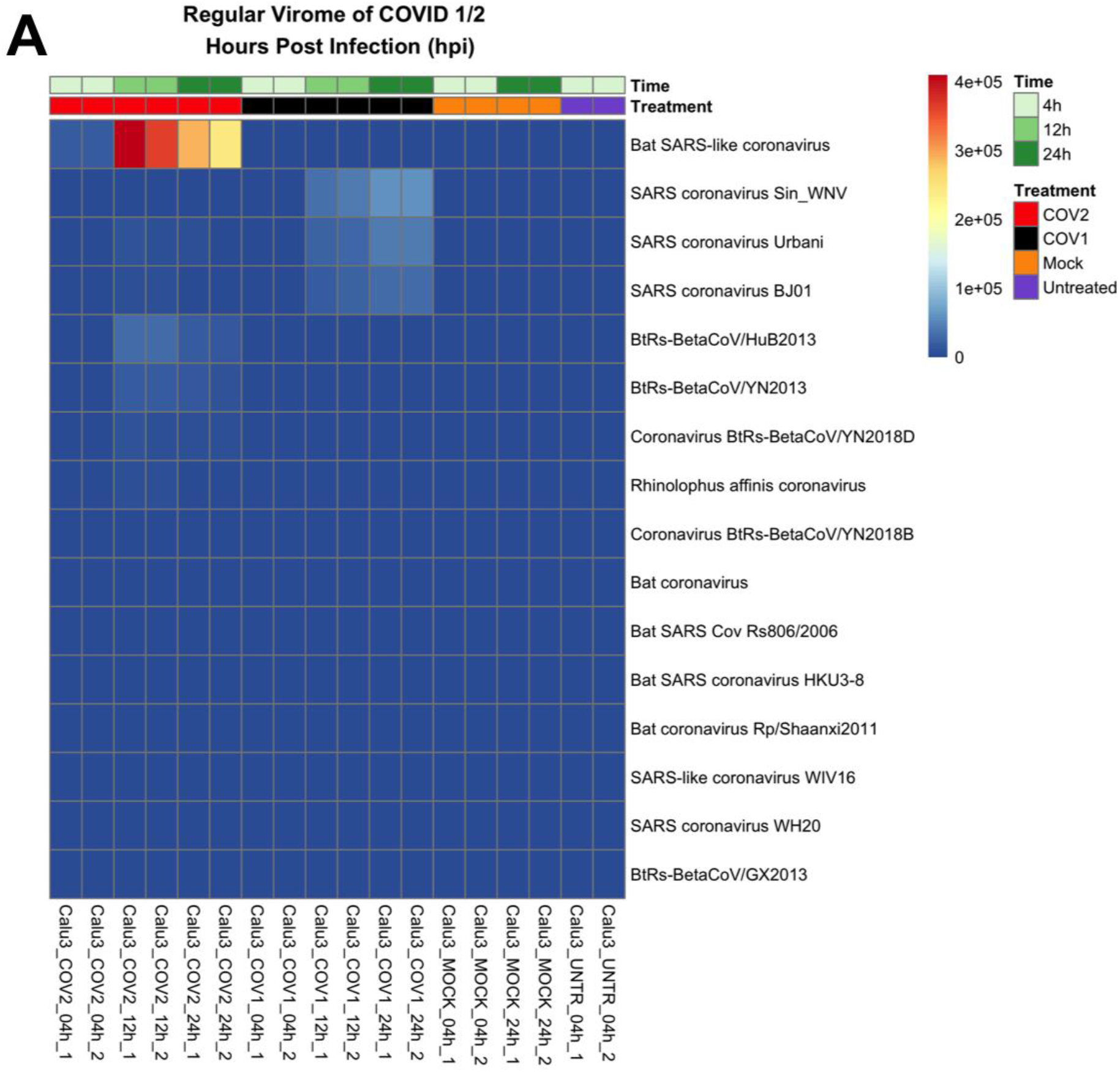
Heatmap of normalized coverage of regular Virome from Emanuel2020 with BLAST to nt database from 05/10/2019. Heatmap of normalized coverage of dark biome contigs binned by species (top 30 species). The nucleotide database was from 5/10/2019 before the discovery of SARS-CoV-2. The top row shows the same row from the main text but identified as a bat SARS like coronavirus.

**Figure S2.**
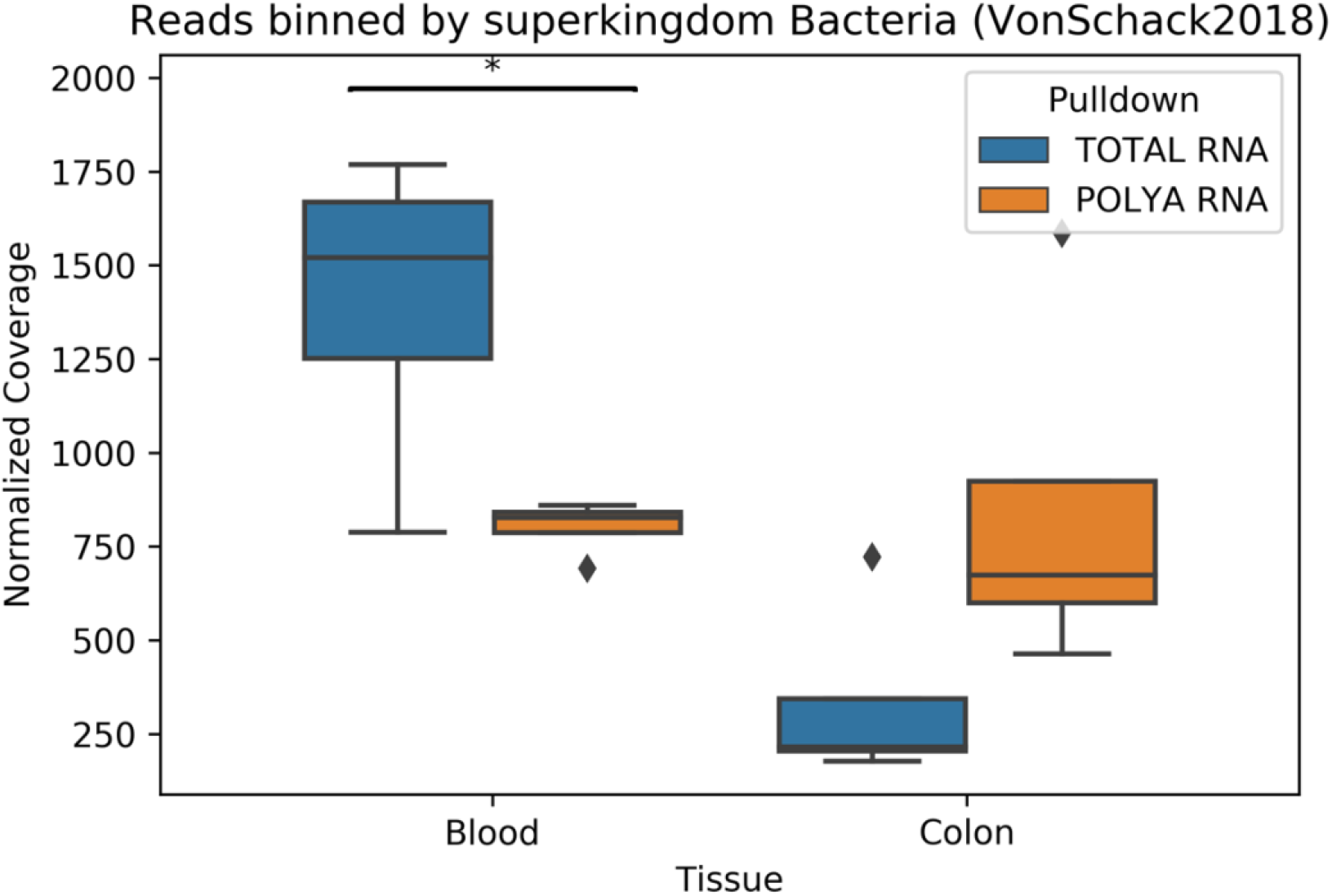
Boxplot of normalized coverage for superkingdom Bacteria in VonSchack2018. Boxplot of normalized coverage of regular biome contigs binned by superkingdom Bacteria. Blood shows significantly more reads in total RNA vs polyA RNA compared to Colon tissue.

**Figure S3.**
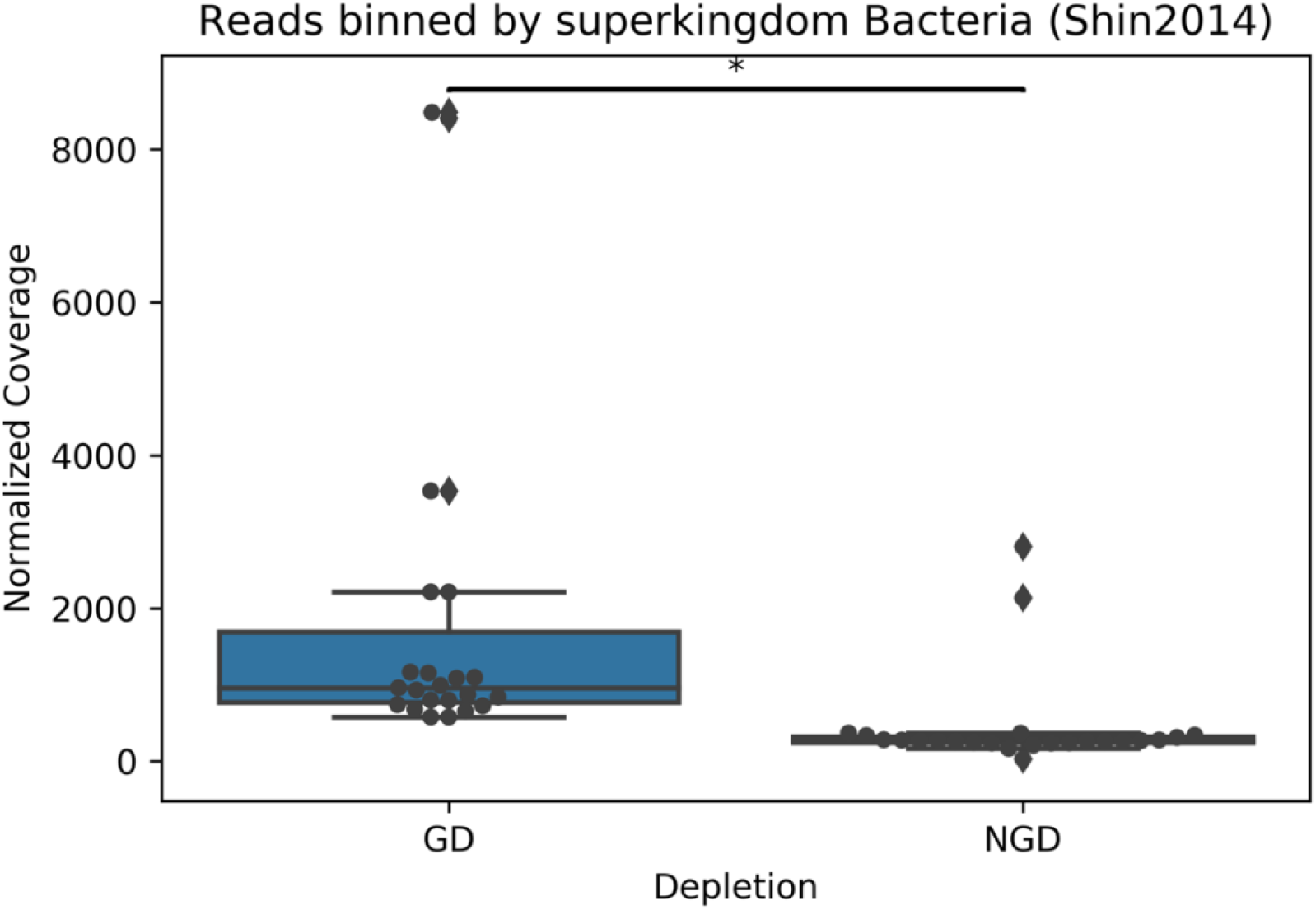
Boxplot of normalized coverage for superkingdom Bacteria in Shin2014. Boxplot of normalized coverage of regular biome contigs binned by superkingdom Bacteria. Globin depletion (GD) has significantly more coverage than non-globin depleted (NGD) blood.

**Figure S4.**
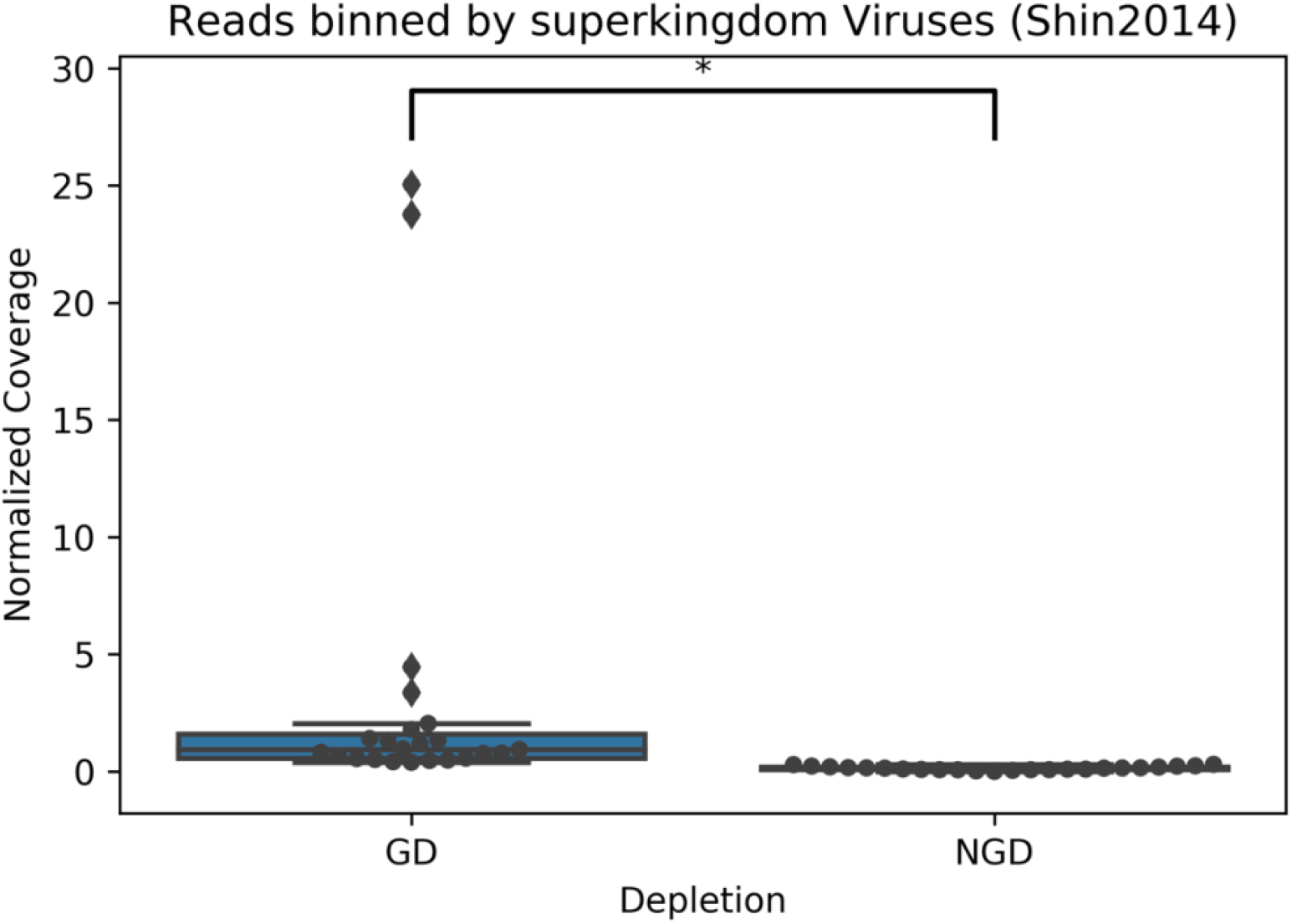
Boxplot of normalized coverage for superkingdom Viruses in Shin2014. Boxplot of normalized coverage of regular biome contigs binned by superkingdom Viruses. Globin depletion (GD) has significantly more coverage than non-globin depleted (NGD) blood.

**Figure S5.**
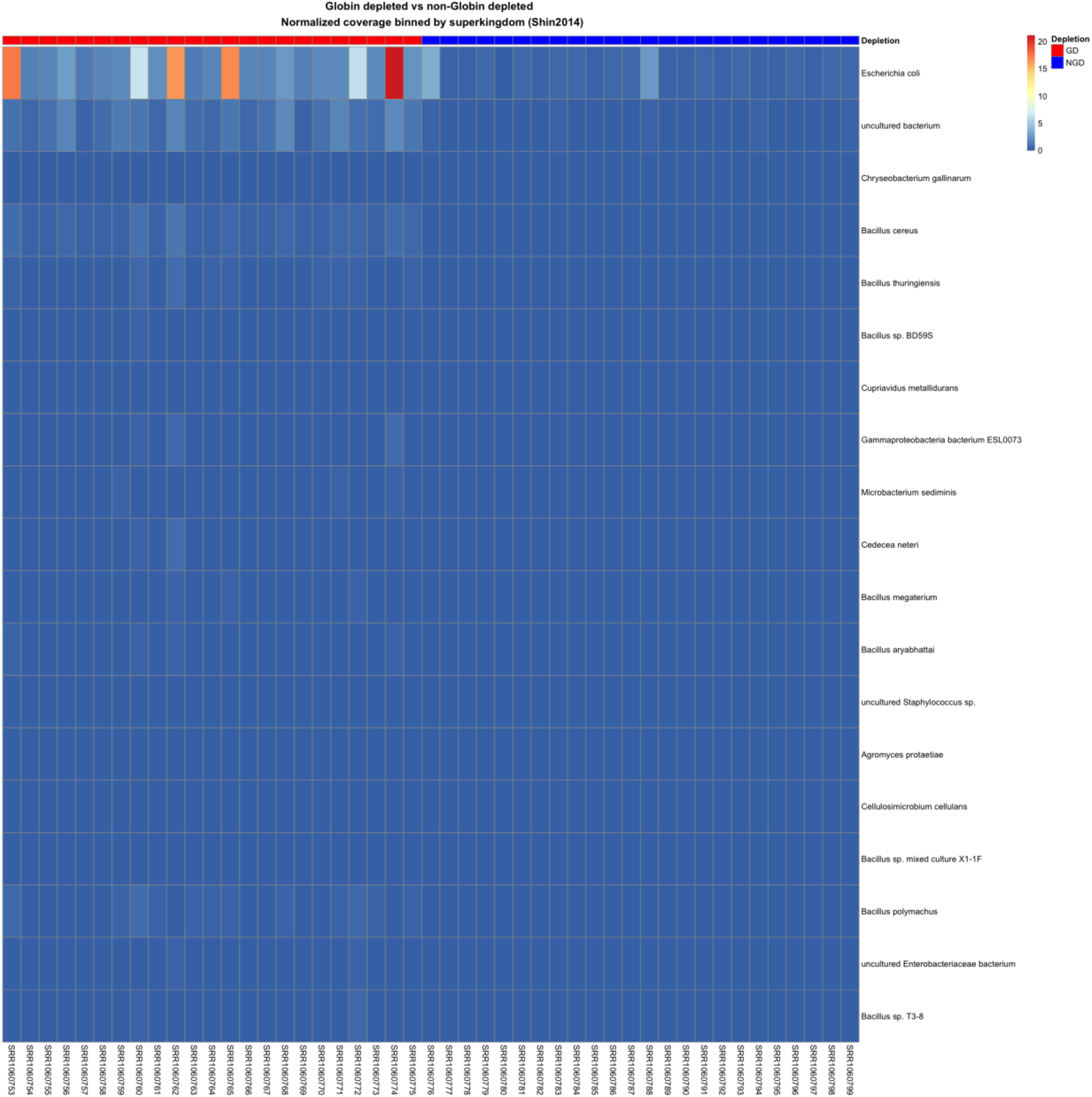
Heatmap of normalized coverage of regular Bacteriome binned by species from Shin2014. Heatmap of normalized coverage of regular biome contigs binned by bacteria species (top 20 species shown for brevity). Globin depletion (GD) is red and non-globin depletion is blue (NGD).

**Figure S5.**
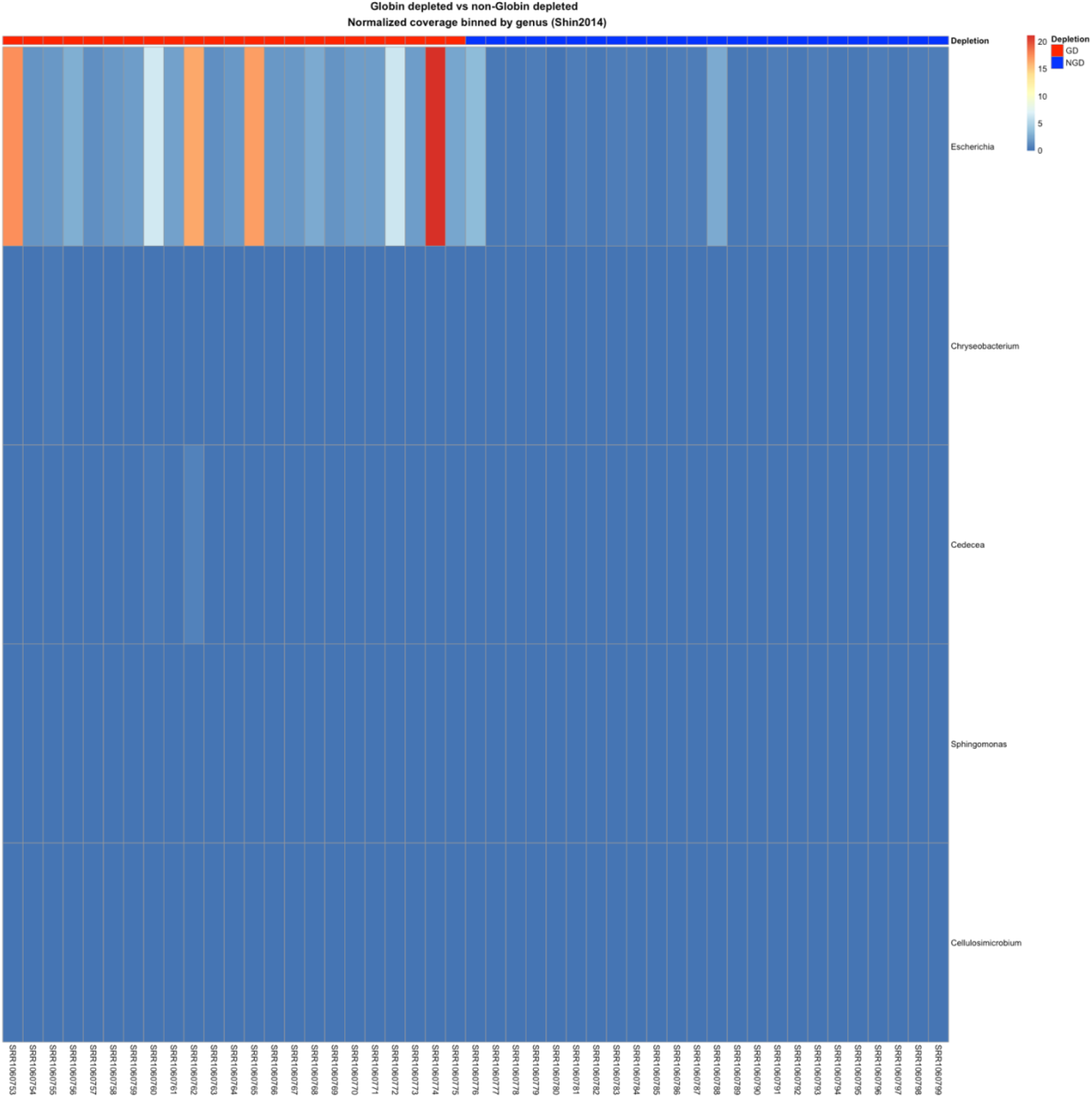
Heatmap of normalized coverage of regular Bacteriome binned by genus from Shin2014. Heatmap of normalized coverage of regular biome contigs binned by bacteria genus. Globin depletion (GD) is red and non-globin depletion is blue (NGD).

**Figure S6.**
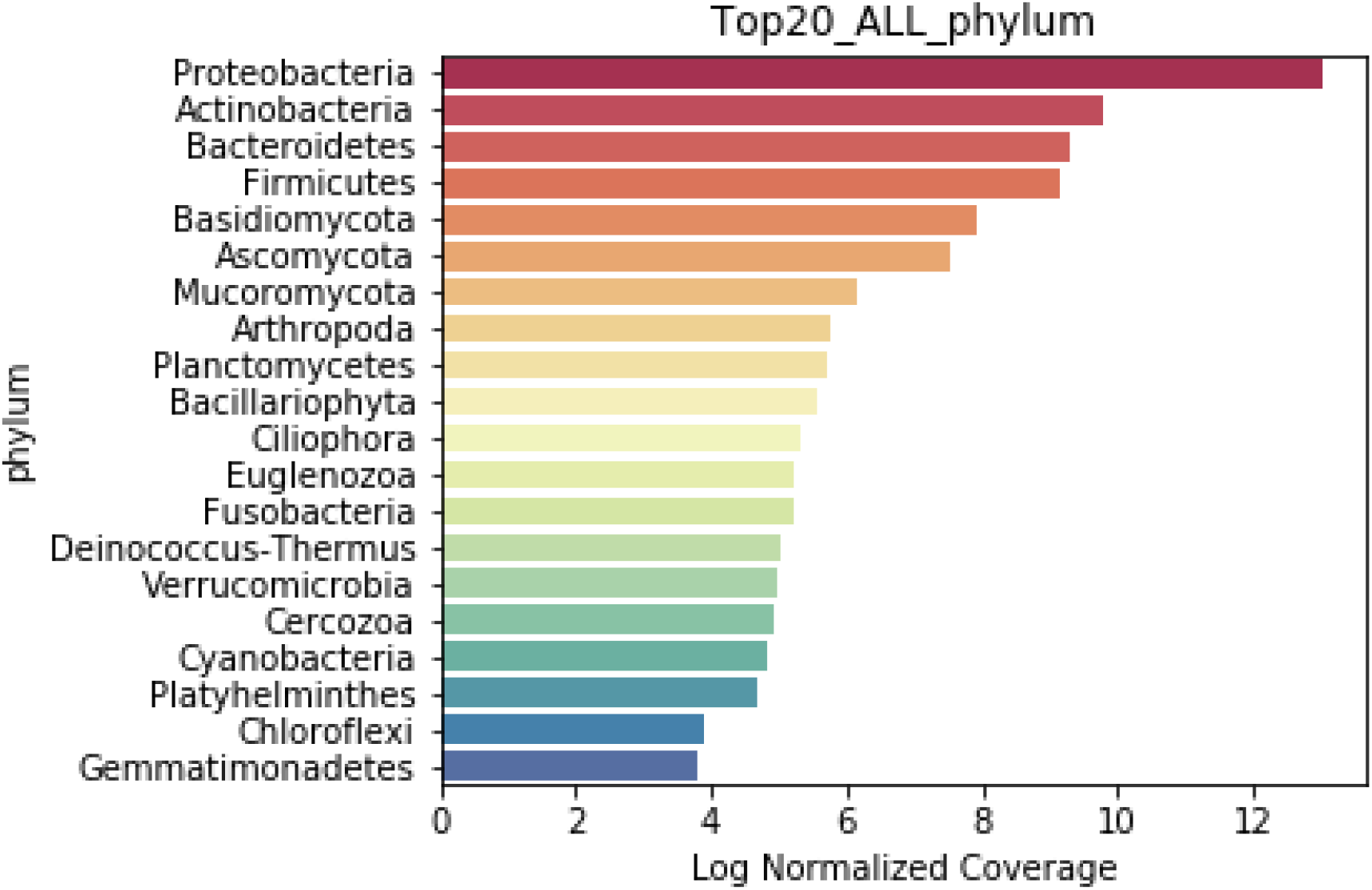
Log coverage binned by phylum from our ALS dataset. Coverage is summed for all of the samples and *alpha proteo-bacteria, Actinobacteria, Firmicutes*, and *Bacteroidetes* are the most highly represented.

**Figure S7.**
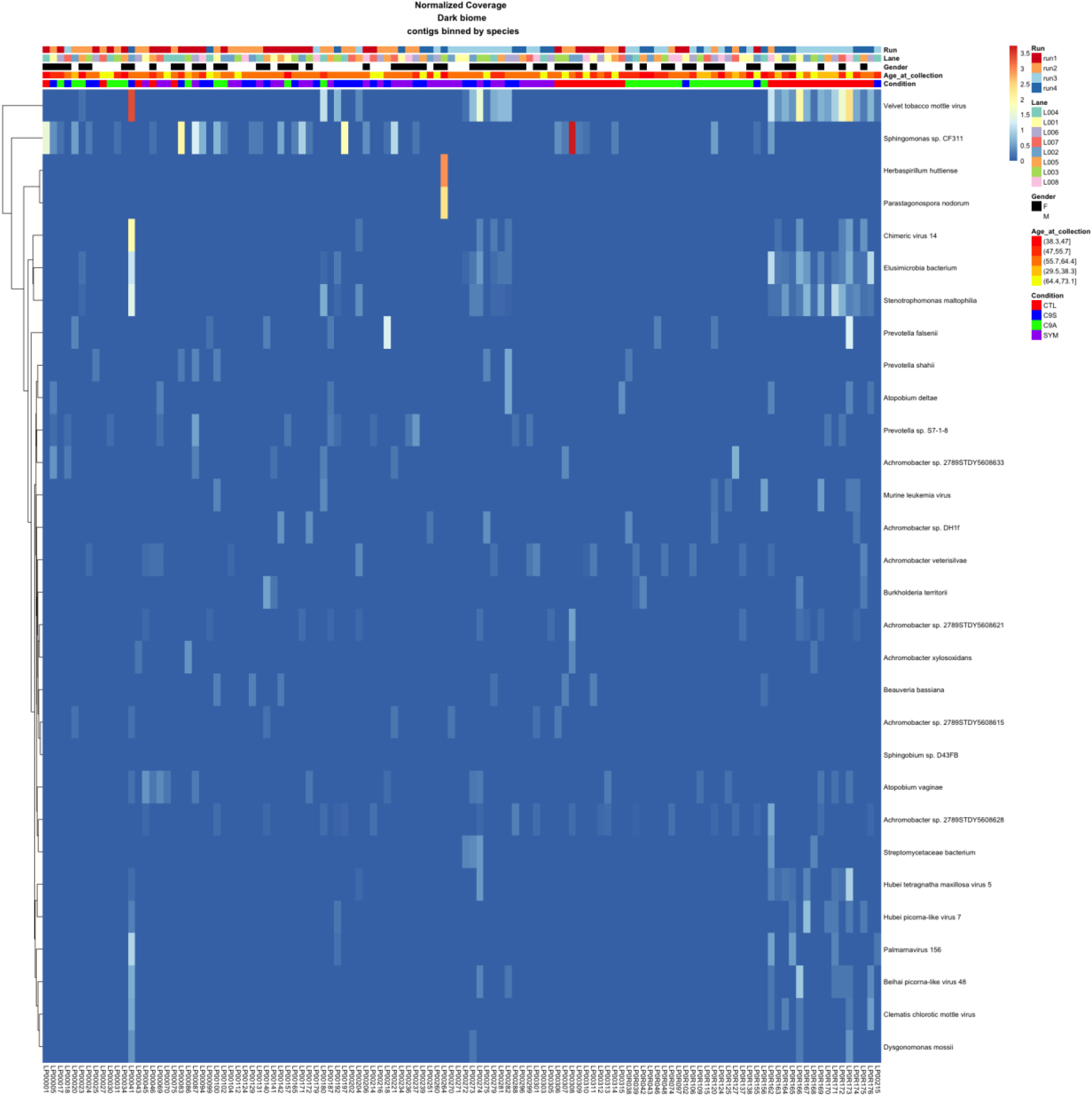
Heatmap of normalized coverage of dark biome contigs binned by species with metadata. Heatmap of normalized coverage of dark biome contigs binned by species (top 30 species shown for brevity). The highest coverage belongs to contigs that show high similarity to velvet tobacco mettle virus. Zero coverage is blue and goes to red with increasing values. These samples were from four conditions including control patients [(CTL) green], ALS symptomatic patients [(SYM) purple], C9-ORF positive ALS symptomatic patients [(C9S) blue] and C9-ORF positive asymptomatic patients [(C9A) red]. Other metadata include gender, lane, run, and age at collection.

**Figure S8.**
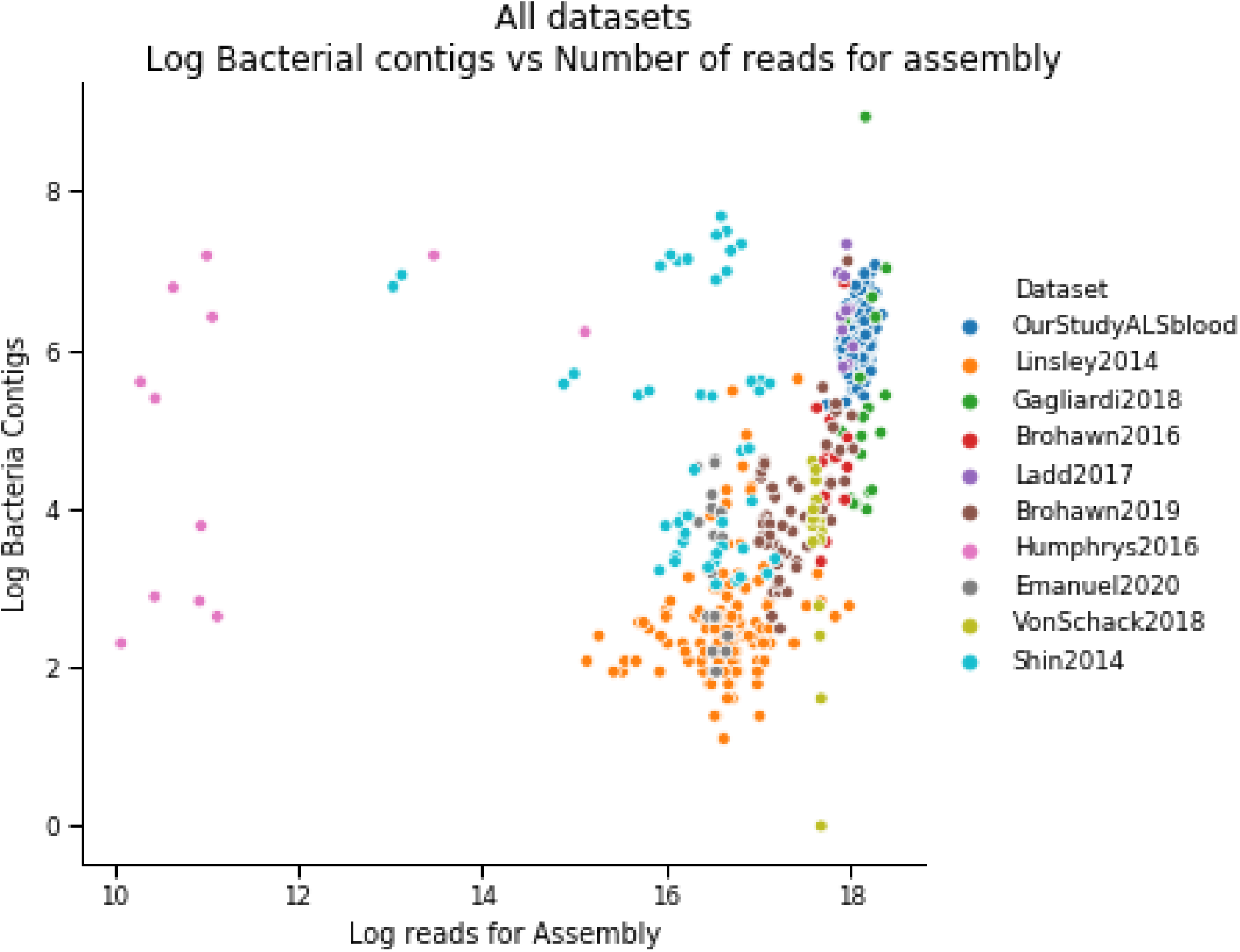
Log Bacterial contigs vs log reads for Assembly. Scatterplot where each dot is a sample from a dataset with log number of Bacterial contigs assembled on the Y-axis and Log reads used in SPAdes on the X-axis. Aside from the Shin, Humphrys, and Emanuel datasets there is a general trend of increased number of bacterial contigs with amount of reads.

**Figure S9.**
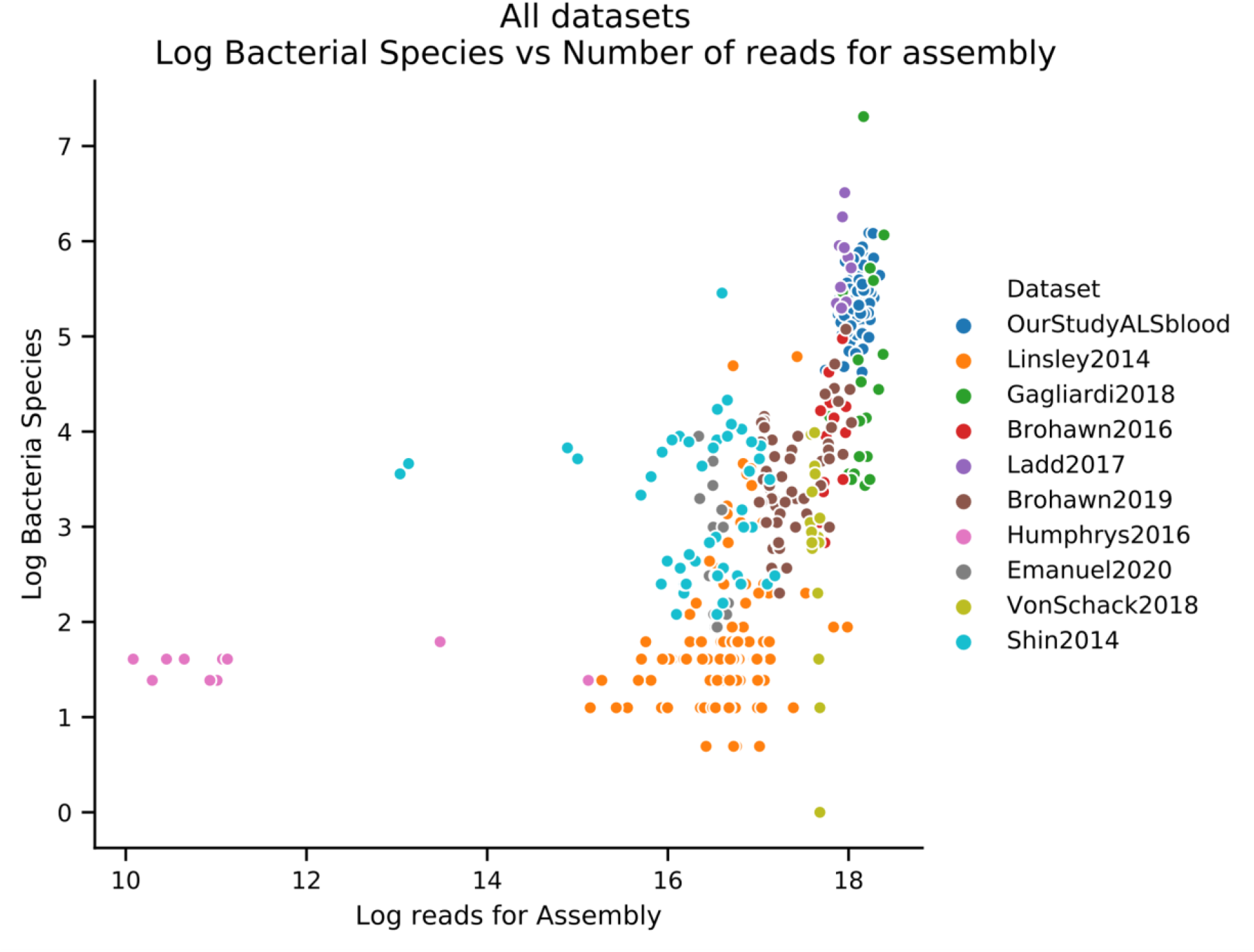
Log number of bacterial species vs log reads for Assembly. Scatterplot where each dot is a sample from a dataset with log number of number of bacterial species detected on the Y-axis and Log reads used in SPAdes on the X-axis.

**Figure S10.**
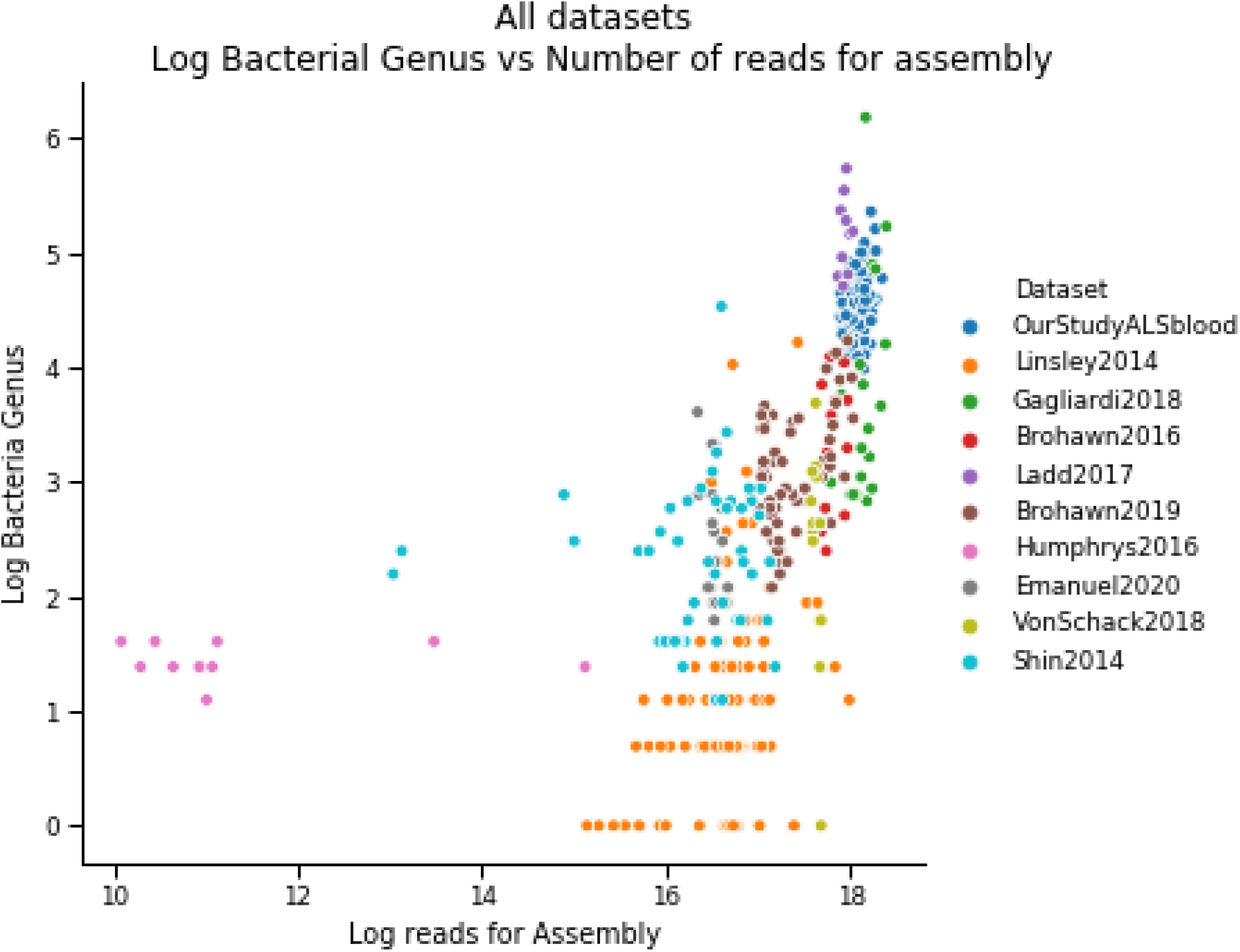
Log number of bacterial genus vs log reads for Assembly. Scatterplot where each dot is a sample from a dataset with log number of number of bacterial genus detected on the Y-axis and Log reads used in SPAdes on the X-axis.

**Figure S11.**
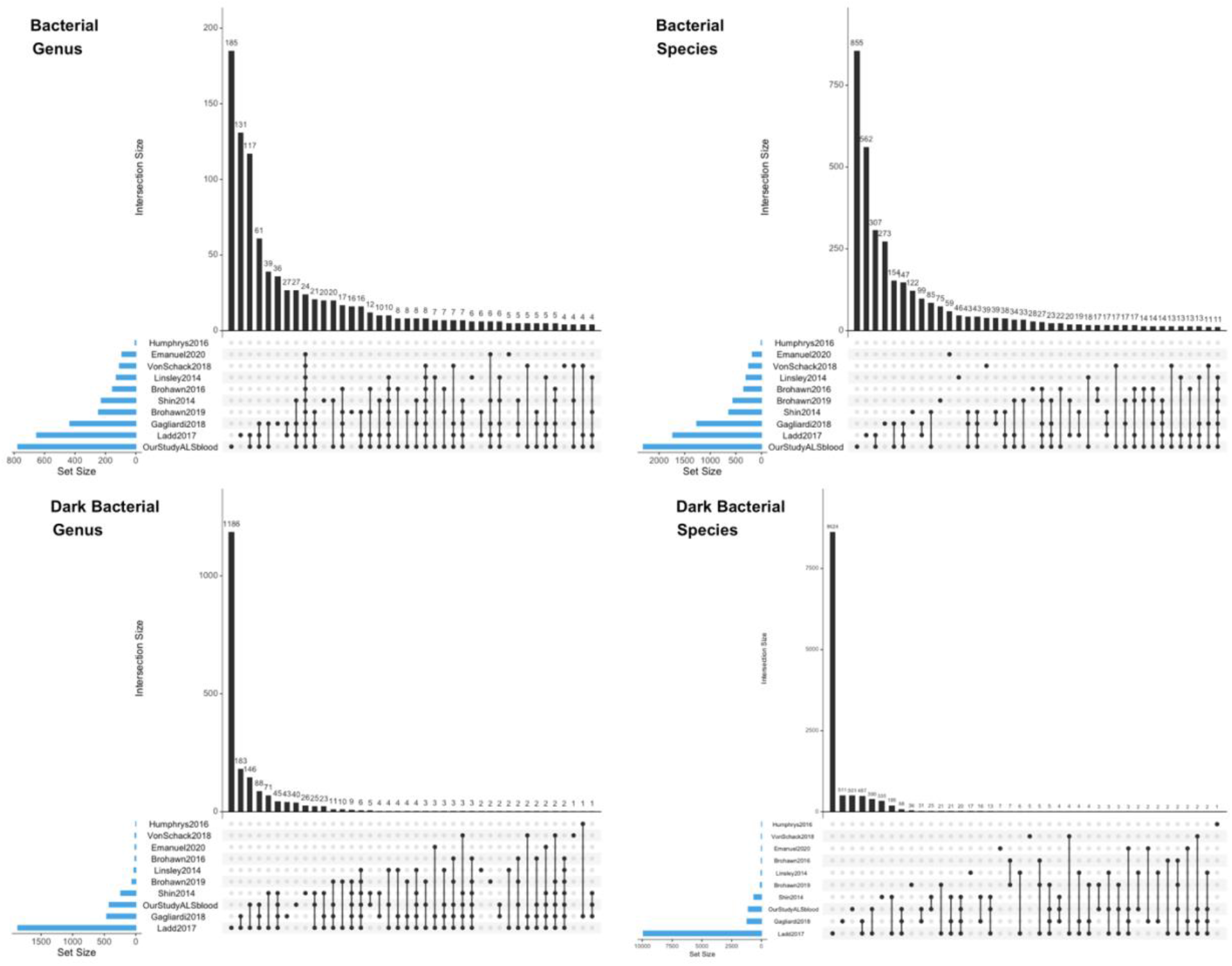
Upset plots of Bacteria for genus/species of regular/dark genome. Upset plots are venn diagram-like plots. Each set is on a row with total amounts in a set as a blue bar plot on the left. The black histogram on top shows the counts that are in the intersection of sets (a single dot for one set or connected dots for multiple sets). The highest number of overlapping bacterial genus is between our dataset and Ladd2017 (133) followed by the intersection between our dataset, Ladd2017 and Gagliardi2018 (61) and there is a modest overlap (24) for 9/10 datasets. This is roughly similar in the Bacterial species figure and in general the larger datasets have more unique and the highest number of overlap.

**Figure S12.**
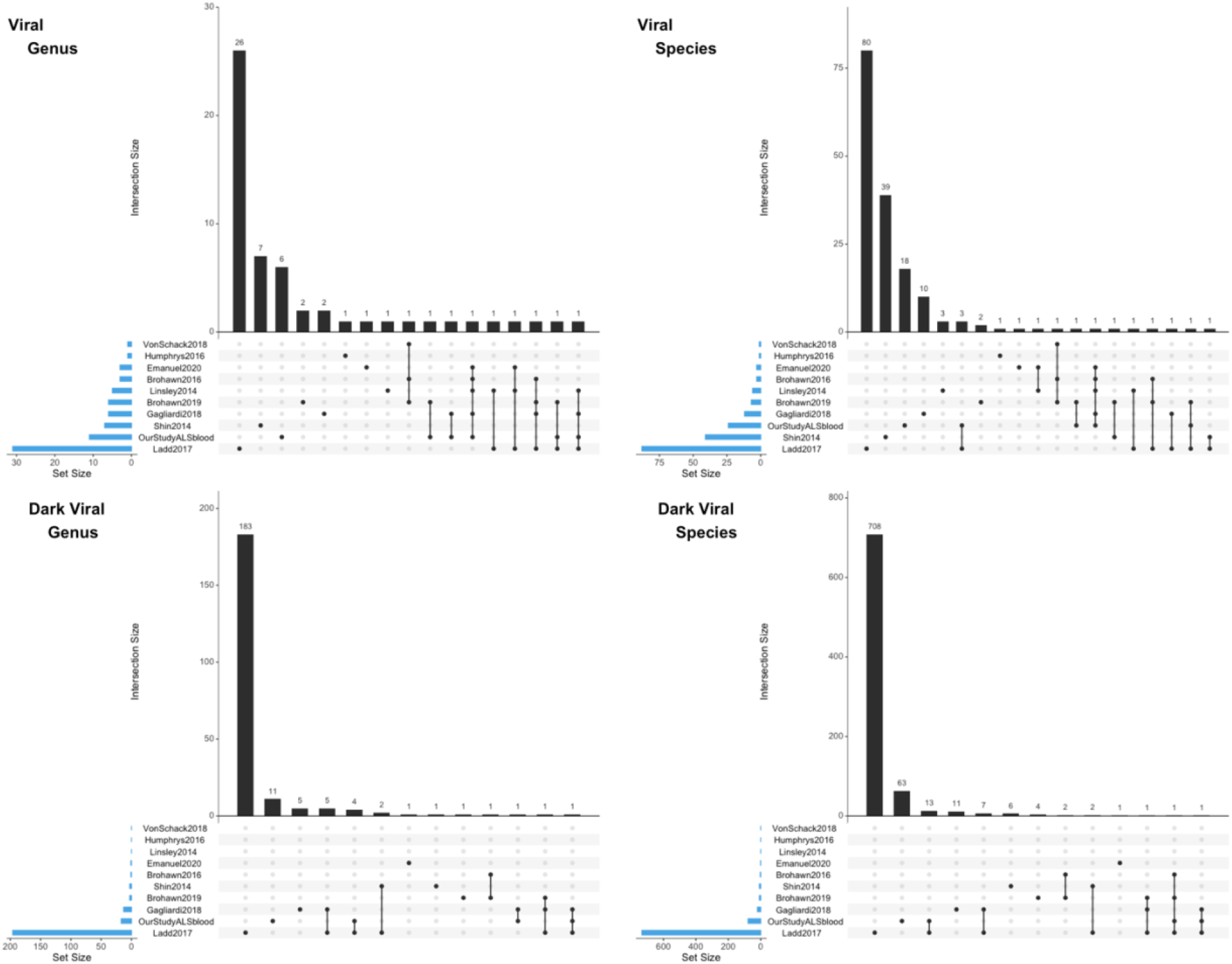
Upset plots of Viruses for genus/species of regular/dark genome. Upset plots are venn diagram-like plots. Each set is on a row with total amounts in a set as a blue bar plot on the left. The black histogram on top shows the counts that are in the intersection of sets (a single dot for one set or connected dots for multiple sets). The regular virome of each dataset is relatively unique with very low amounts of overlap (<= 3) between datasets (species and genus shows a similar pattern). Interestingly, the highest overlap for species in the dark virome is between our dataset and Ladd2017 (13).

**Figure S13.**
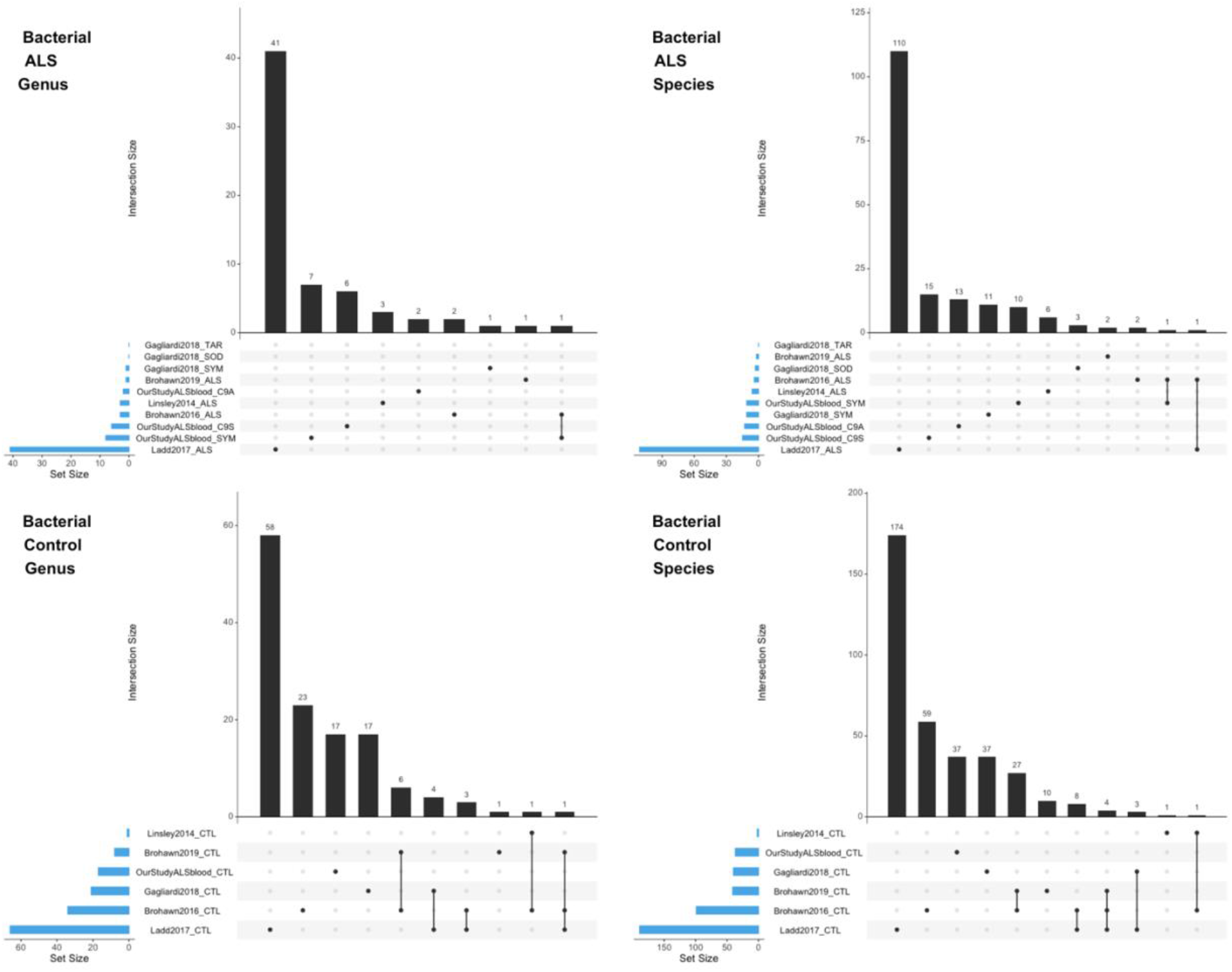
Upset plots of Bacteria in the regular biome for genus/species in ALS and Control contigs. Upset plots are venn diagram-like plots. Each set is on a row with total amounts in a set as a blue bar plot on the left. The black histogram on top shows the counts that are in the intersection of sets (a single dot for one set or connected dots for multiple sets). We assigned a contig to a condition if >= 2 samples from that condition contain at least 90% of the summed normalized coverage (from all samples) to the contig. In the genus and species from ALS samples there is a low amount of overlap between datasets (<= 1). When we look at control samples there is a much higher overlap for both genus and species.

**Figure S13.**
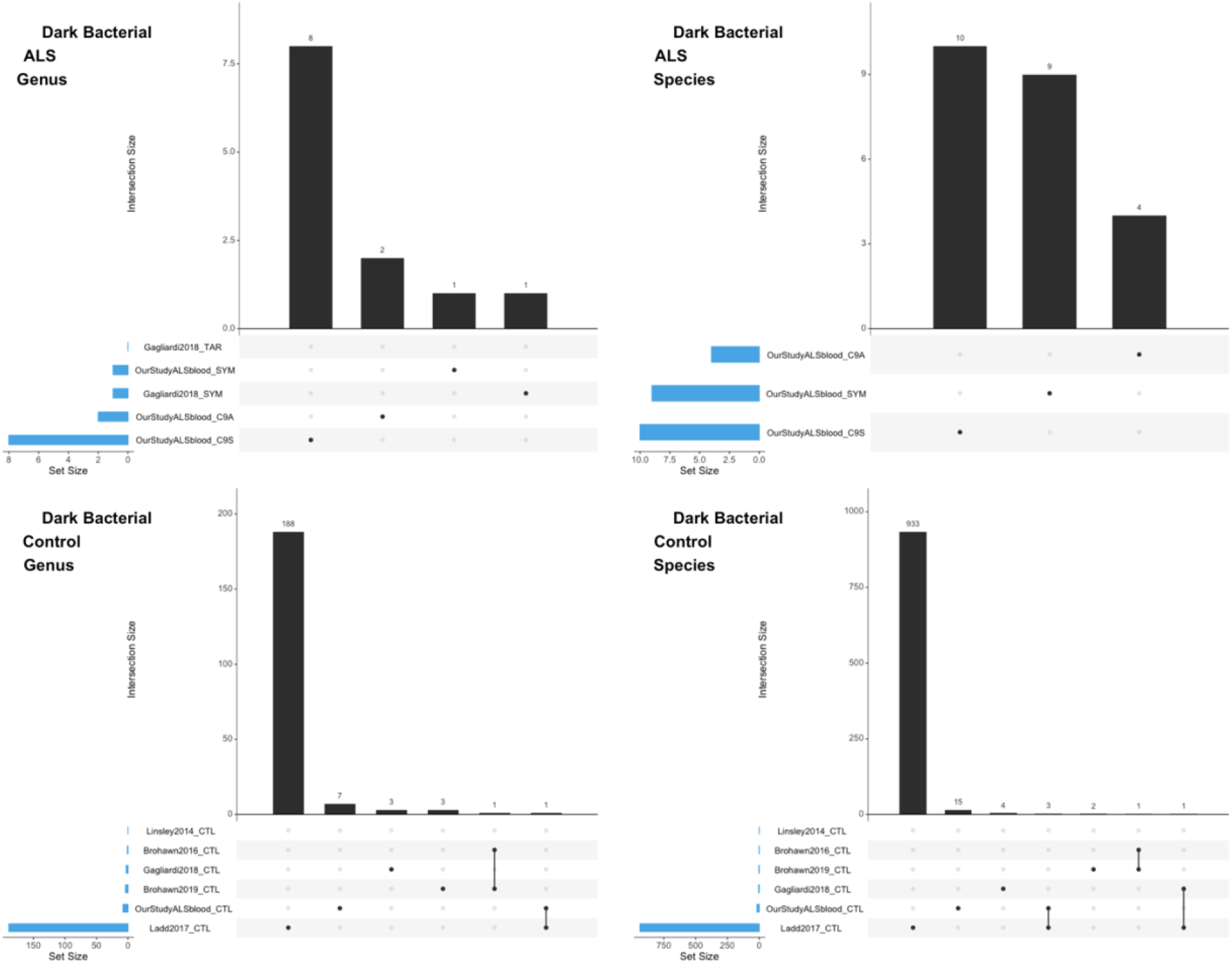
Upset plots of Bacteria in the dark biome for genus/species in ALS and Control contigs. Upset plots are venn diagram-like plots. Each set is on a row with total amounts in a set as a blue bar plot on the left. The black histogram on top shows the counts that are in the intersection of sets (a single dot for one set or connected dots for multiple sets). We assigned a contig to a condition if >= 2 samples from that condition contain at least 90% of the summed normalized coverage (from all samples) to the contig. Conditions with no recovered viruses have been omitted for clarity. Similarly to the regular bacteriome, there is no overlap in ALS samples and a small amount of overlap in the conditions.

